# Ecological divergence in sympatry causes gene misregulation in hybrids

**DOI:** 10.1101/717025

**Authors:** Joseph A. McGirr, Christopher H. Martin

## Abstract

Ecological speciation occurs when reproductive isolation evolves as a byproduct of adaptive divergence between populations. However, it is unknown whether divergent ecological selection on gene regulation can directly cause reproductive isolation. Selection favoring regulatory divergence between species could result in gene misregulation in F1 hybrids and ultimately lower hybrid fitness. We combined 58 resequenced genomes with 124 transcriptomes to test this hypothesis in a young, sympatric radiation of *Cyprinodon* pupfishes endemic to San Salvador Island, Bahamas, which consists of a dietary generalist and two novel trophic specialists – a molluscivore and a scale-eater. We found more differential gene expression between closely related sympatric specialists than between allopatric generalist populations separated by 1000 km. Intriguingly, 9.6% of genes that were differentially expressed between sympatric species were also misregulated in their F1 hybrids. Consistent with divergent ecological selection causing misregulation, a subset of these genes were in highly differentiated genomic regions and enriched for functions important for trophic specialization, including head, muscle, and brain development. These regions also included genes that showed evidence of hard selective sweeps and were significantly associated with oral jaw length – the most rapidly diversifying skeletal trait in this radiation. Our results indicate that divergent ecological selection in sympatry can cause hybrid gene misregulation which may act as a primary reproductive barrier between nascent species.

**Significance:** It is unknown whether the same genes that regulate ecological traits can simultaneously contribute to reproductive barriers between species. We measured gene expression in two trophic specialist species of *Cyprinodon* pupfishes that rapidly diverged from a generalist ancestor. We found genes differentially expressed between species that also showed extreme expression levels in their hybrid offspring. Many of these genes showed signs of selection and have putative effects on the development of traits that are important for ecological specialization. This suggests that genetic variants contributing to adaptive trait divergence between parental species negatively interact to cause hybrid gene misregulation, potentially producing unfit hybrids. Such loci may be important barriers to gene flow during the early stages of speciation, even in sympatry.

## Introduction

Adaptive radiations showcase dramatic instances of biological diversification resulting from ecological speciation, which occurs when reproductive isolation (RI) evolves as a byproduct of adaptive divergence between populations (1, 2). Ecological speciation predicts that populations adapting to different niches will accumulate genetic differences due to divergent ecological selection, indirectly resulting in reduced gene flow. Gene regulation is a major target of selection during adaptive divergence, with many known cases of divergent gene regulation underlying ecological traits (3–7). However, it is still unknown whether divergent ecological selection on gene regulation contributes to reproductive barriers during speciation (8, 9).

Hybridization between ecologically divergent populations can break up coadapted genetic variation, resulting in (Bateson) Dobzhansky-Muller incompatibilities (DMIs) if divergent alleles from parental populations are incompatible in hybrids and cause reduced fitness (10, 11). DMIs can result in gene misregulation: transgressive expression levels that are significantly higher or lower in F1 hybrids than either parental population. Because gene expression is largely constrained by stabilizing selection, gene misregulation in hybrids is expected to disrupt highly coordinated developmental processes and reduce fitness (12, 13). Indeed, crosses between distantly related species show that misregulation is often associated with reduced hybrid fitness in the form of hybrid sterility and inviability (i.e. intrinsic postzygotic isolation) (14–16). DMIs causing these forms of strong intrinsic isolation evolve more slowly than premating isolating barriers and are traditionally modeled as fixed genetic variation between allopatric populations (11).

However, it is unknown whether hybrid gene misregulation also contributes to RI during the early stages of speciation, particularly for populations diverging in sympatry (9, 17, 18). Either segregating or fixed alleles causing gene misregulation in hybrids could disrupt developmental processes resulting in genetic incompatibilities (intrinsic postzygotic isolation) or reduced performance under natural conditions (extrinsic postzygotic isolation). Emerging evidence suggests that weak intrinsic DMIs segregate within natural populations (19) and are abundant between recently diverged species, reaching hundreds of incompatibility loci within swordtail fish hybrid zones (20, 21). Furthermore, hybrid gene misregulation has been reported at early stages of divergence within a species of intertidal copepod (22) and between young species of lake whitefish (23).

We hypothesized that regulatory genetic variants causing adaptive expression divergence between sympatric species may negatively interact to cause misregulation and reduced fitness in hybrids. Such incompatible alleles could promote rapid speciation because they would simultaneously contribute to adaptive trait divergence and reduce gene flow between populations (18, 24, 25). Here we tested this hypothesis in a young (10 kya), sympatric radiation of *Cyprinodon* pupfishes endemic to San Salvador Island, Bahamas. This radiation consists of a dietary generalist and two derived specialists adapted to novel trophic niches: a molluscivore (*C. brontotheroides*) and a scale-eater (*C. desquamator*) (26). Hybrids among these species exhibit reduced fitness in the wild and impaired feeding performance in the lab (27, 28). We took a genome-wide approach to identify genetic variation underlying F1 hybrid gene misregulation and found 125 ecological DMI candidate genes that were misregulated, highly differentiated between populations, and strikingly enriched for developmental functions related to trophic specialization. Our findings show that regulatory variation underlying adaptive changes in gene expression can interact to cause hybrid gene misregulation, which may contribute to reduced hybrid fitness and restrict gene flow between sympatric populations.

## Results

### Trophic specialization, not geographic distance, drives major changes in gene expression and hybrid gene misregulation

We sampled two lake populations on San Salvador Island (Crescent Pond and Osprey Lake) in which generalist pupfish coexist with the endemic molluscivore and scale-eater specialist species. We also collected outgroup generalist populations from North Carolina, USA and New Providence Island, Bahamas (Fig. 1A). Wild caught fishes and their F1 offspring were reared in a common laboratory environment. Overall, genetic divergence increased with geographic distance between allopatric generalist populations and was lowest between sympatric populations (Table S1; genome-wide mean *F_st_* measured across 13.8 million SNPs: San Salvador generalists vs. North Carolina = 0.217; vs. New Providence = 0.155; vs. scale-eaters = 0.106; vs. molluscivores = 0.056). We tested whether isolation by distance explained patterns of gene expression divergence and hybrid gene misregulation while controlling for phylogenetic relatedness using a maximum likelihood tree estimated with RAxML from 1.7 million SNPs (Fig. 1; Fig. S1). Geographic distance among populations was a significant predictor of the proportion of differential gene expression between populations at two days post fertilization (2 dpf) (Fig. 1B; phylogenetic generalized least squares (PGLS); *P* = 0.02). This is consistent with a model of gene expression evolution governed largely by stabilizing selection and drift (29, 30). However, at eight days post fertilization (8 dpf), when craniofacial structures of the skull begin to ossify (31), geographic distance was no longer associated with differential expression (Fig. 1C; PGLS; *P* = 0.18), which was higher between sympatric trophic specialist species on San Salvador Island than between generalist populations spanning 1000 km across the Caribbean.

**Fig. 1.**
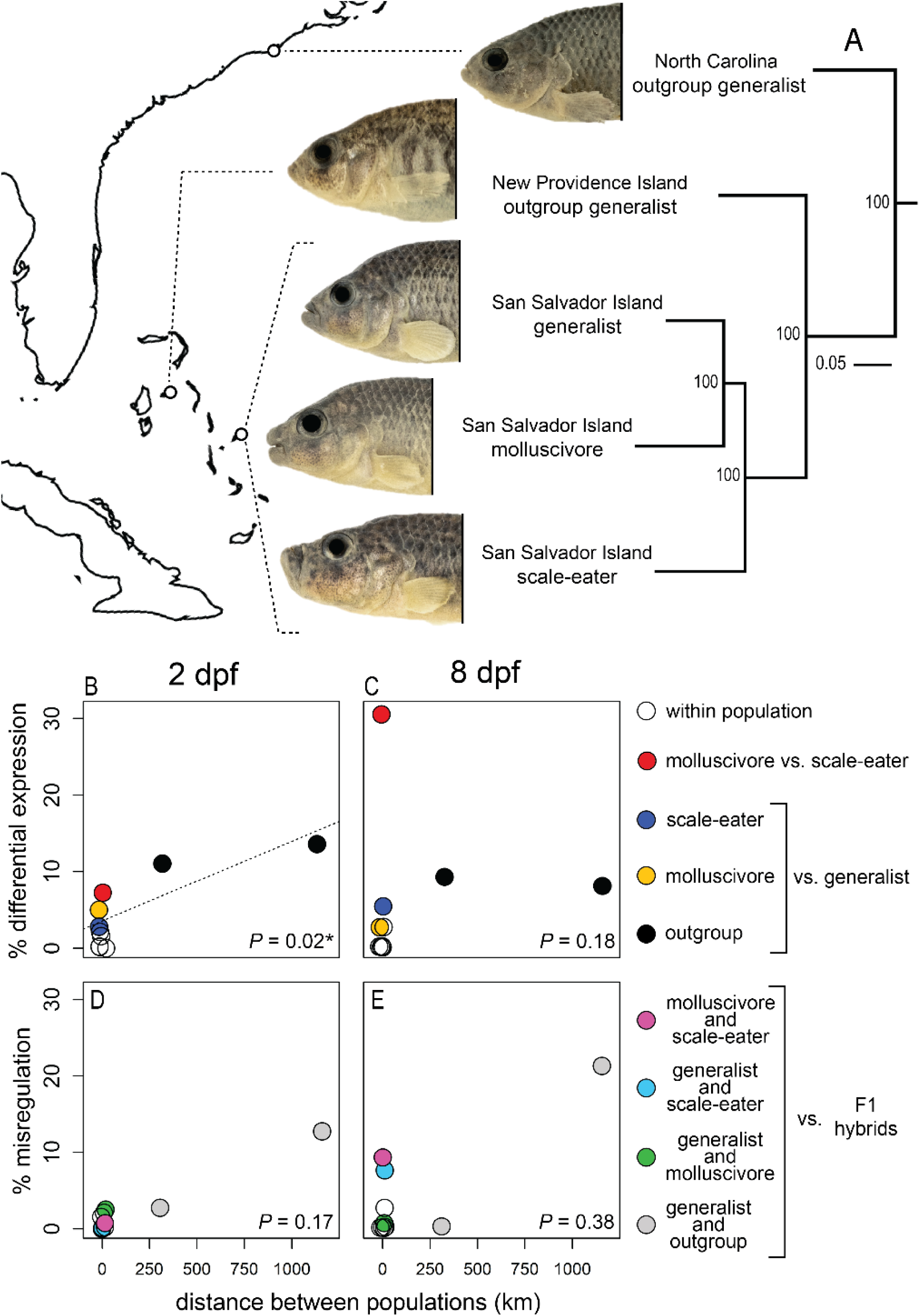
Caribbean-wide patterns of gene expression and misregulation across sympatric and allopatric populations of *Cyprinodon* pupfishes. A) Maximum likelihood tree estimated from 1.7 million SNPs showing phylogenetic relationships among generalist populations and specialist species (100% bootstrap support indicated at nodes). B) Geographic distance separating populations was associated with differential gene expression levels in embryos at 2 days post fertilization (2 dpf; phylogenetic least squares *P* = 0.02, dotted regression line). C) In whole larvae at 8 dpf differential expression was not associated with geographic distance (PGLS; *P* = 0.18) and was higher between sympatric specialists (red) than between allopatric generalists separated by 300 and 1000 km (black). D and E) Hybrid misregulation for sympatric crosses at 8 dpf than 2 dpf. Geographic distance was not associated with hybrid misregulation at either developmental stage (PGLS; 2 dpf *P* = 0.17; 8dpf *P* = 0.38). Percentages in B-E were measured using Crescent Pond crosses.

Geographic distance between parental populations was not associated with gene misregulation in F1 hybrids at either developmental stage (Fig. 1D and E; PGLS; 2 dpf *P* = 0.17; 8dpf *P* = 0.38). 9.3% of genes were misregulated in specialist F1 hybrids (Fig. 1E; Crescent Pond molluscivore × scale-eater), comparable to species pairs with much greater divergence times (16, 32). Out of 3,669 misregulated genes containing heterozygous sites in F1 hybrids that were homozygous in parents, 819 (22.3%) showed allele specific expression and were not differentially expressed between parental populations – patterns consistent with compensatory regulation underlying misregulation (Fig. S2-4, Table S2).

### Genes showing divergent expression between species are also misregulated in their F1 hybrids

We used two approaches to identify gene misregulation associated with ecological divergence between species. First, we found 716 genes that showed differential expression between San Salvador species that were also misregulated in their F1 hybrids (Fig. 2, Table S3). Nearly all these genes (99.4%) were misregulated in only one lake population and 69.8% were only misregulated at 8 dpf in comparisons involving scale-eaters (Fig. 2A-H). Four genes showed differential expression between species and hybrid misregulation in both lake comparisons (*trim47*, *krt13*, *s100a1*, *elovl7*; Table S4).

**Fig. 2.**
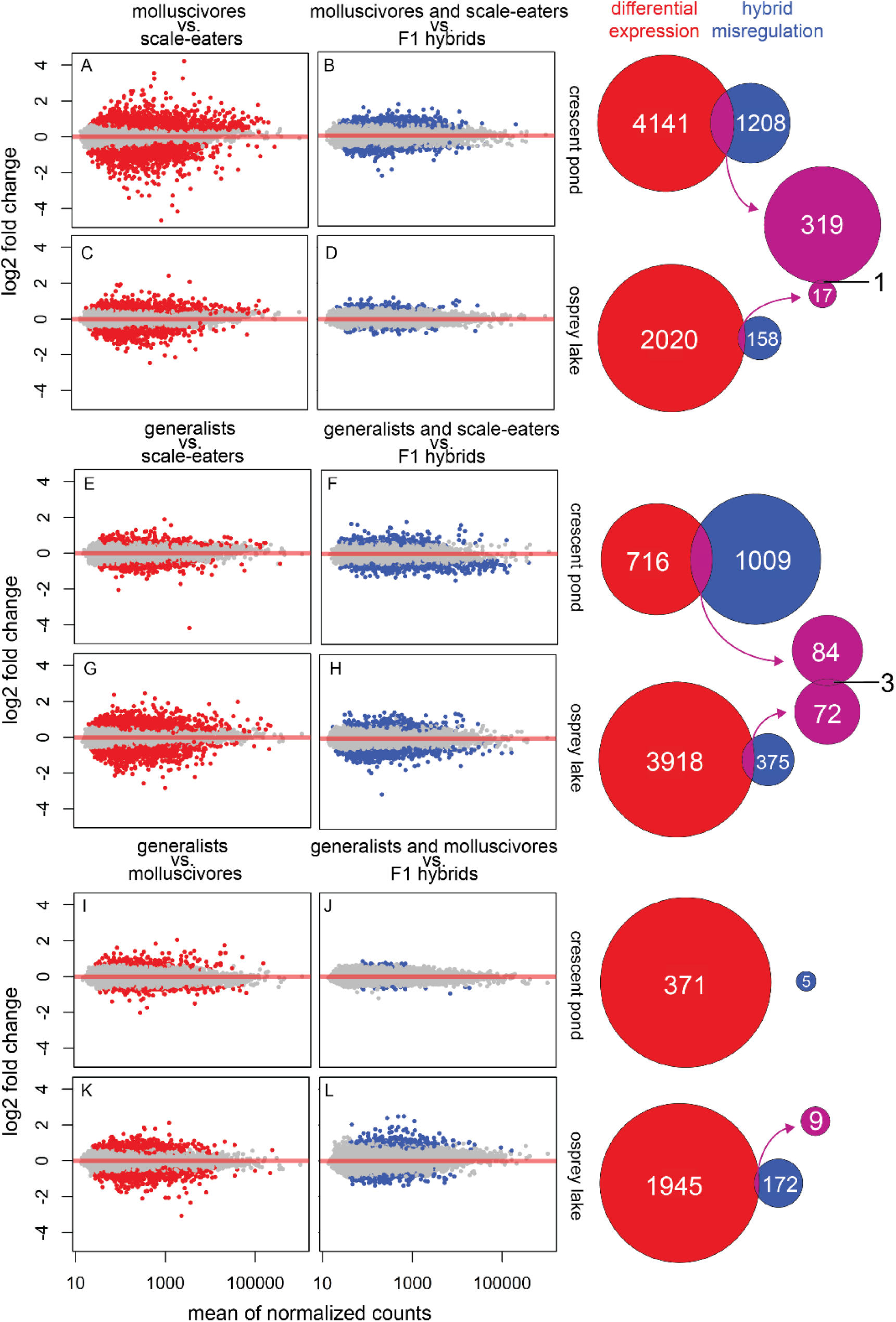
Genes differentially expressed between species are misregulated in their F1 hybrids at 8 days post fertilization. Genes differentially expressed between San Salvador species from Crescent Pond and Osprey Lake are shown in red for molluscivore × scale-eater crosses (A-D), generalist × scale-eater crosses (E-H), and generalist × molluscivore crosses (I-L). Genes misregulated in F1 hybrids are shown in blue. In comparisons involving reciprocal crosses (D, J, and L), we only show genes misregulated in a single cross direction. A total of 716 genes (purple) were differentially expressed between species and also misregulated in their F1 hybrids. Purple Venn diagrams show overlap between lake population comparisons; 4 genes showed differential expression and misregulation in both lake comparisons.

Second, we identified genes showing parallel expression divergence in both specialist species relative to generalists that were misregulated in specialist F1 hybrids (Fig. 3). This pattern likely results from parallel expression in molluscivores and scale-eaters controlled by different genetic mechanisms (33). Significantly more genes showed differential expression in both specialist comparisons than expected by chance (Fig. 3A-D; Fisher’s exact test, *P* < 2.7 × 10^-5^). Of these, 96.6% (1,206) showed the same direction of expression in specialists relative to generalists, which was more than expected under a neutral model of gene expression evolution (Fig. 3E and F; binomial test, *P* < 1.0 × 10^-16^). 45 of the 1,206 genes showing parallel expression divergence in specialists also showed misregulation in specialist F1 hybrids (Fig. 3F). Eight of these genes were severely misregulated to the extent that they were differentially expressed in hybrids relative to all other populations in our dataset. For example, *sypl1* showed significantly higher expression in 8 dpf Crescent Pond molluscivore × scale-eater F1 hybrids than all other crosses spanning 1000 km from San Salvador Island, Bahamas to North Carolina, USA (*P* = 2.35 × 10^-4^; Fig. 3G). Overexpression of this gene is associated with epithelial-mesenchymal transition, an important process during cranial neural crest cell migration (34, 35). Similarly, *scn4a* showed significantly lower expression in 8 dpf Crescent Pond specialist F1 hybrids than all other crosses (*P* = 5.49 × 10^-4^; Fig. 3H). Mutations in this gene are known to cause paramyotonia congenita, a disorder causing weakness and stiffness of craniofacial skeletal muscles (36).

**Fig. 3.**
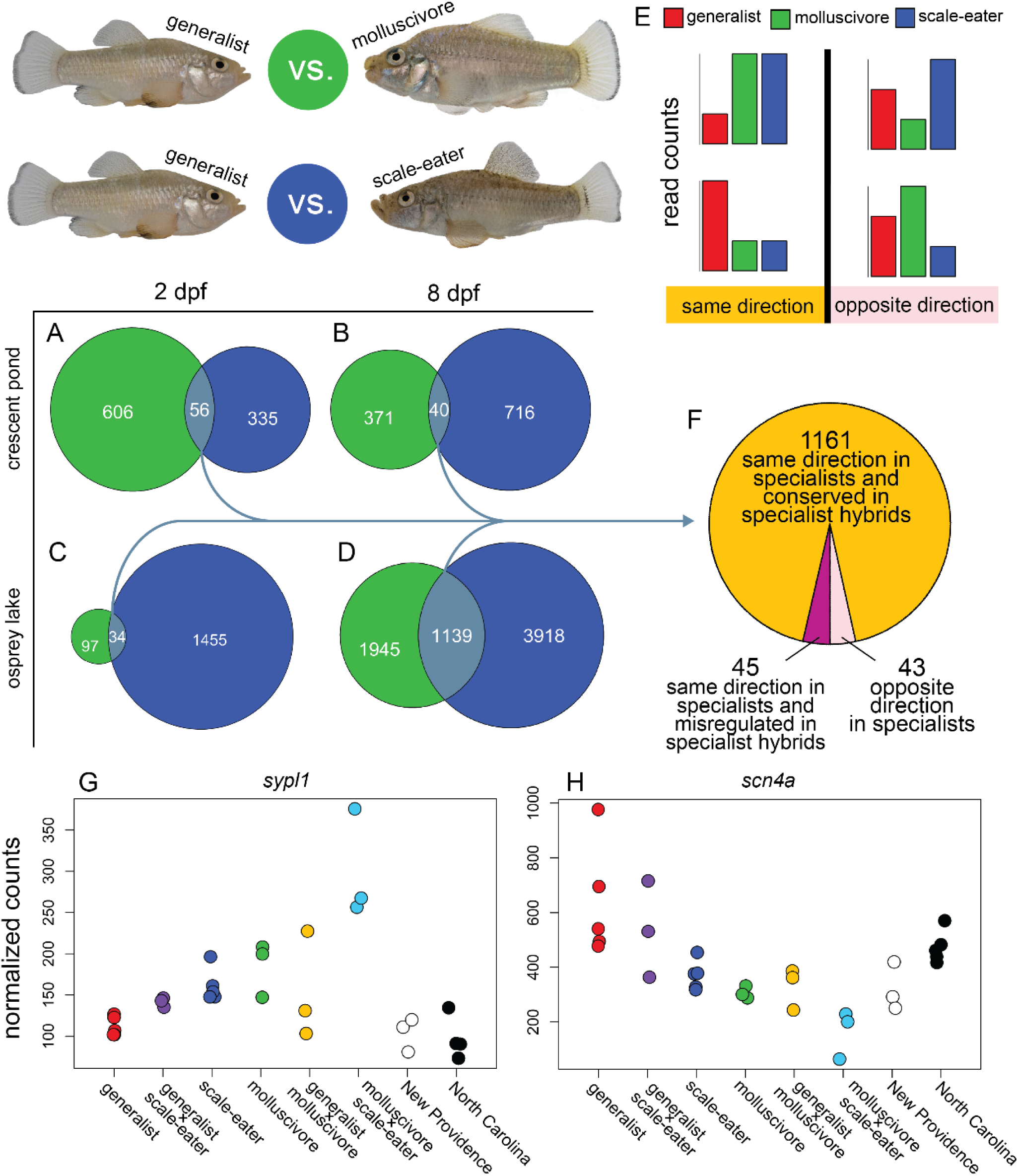
Genes showing parallel expression divergence in specialists are misregulated in specialist hybrids. Genes differentially expressed between generalists and molluscivores (green) were compared to the set of genes differentially expressed between generalists and scale-eaters (dark blue). A-D) Significantly more genes showed differential expression in both specialist comparisons (light blue) than expected by chance in both lakes at both developmental stages (Fisher’s exact test, P < 2.7 × 10^-5^). E) A neutral model of gene expression evolution would predict that only 50% of genes should show the same direction of expression in specialists relative to generalists (yellow). F) Instead, 96.6% of genes showed the same direction of expression in specialists, suggesting significant parallel expression divergence in specialists (Binomial exact test; *P* < 1.0 × 10^-16^). Consistent with incompatible regulatory mechanisms underlying parallel expression in specialists, 45 of these genes were misregulated in specialist F1 hybrids, including G) *sypl1* and H) *scn4a* which showed extreme misregulation: expression levels outside the range of all other Caribbean populations examined.

### Misregulated genes under selection influence adaptive ecological traits in trophic specialists

Out of 750 total unique genes identified above as differentially expressed between populations and misregulated in F1 hybrids, 125 (17%) were within 20 kb of SNPs that were fixed between populations (*F_st_* = 1) and within 20 kb windows showing high absolute genetic divergence between populations (*D_xy_* ≥ genome-wide 90^th^ percentile; range: 0.0031 – 0.0075; Table S1). This set of 125 genes, which we refer to as ecological DMI candidate genes, was significantly enriched for functional categories highly relevant to divergent specialist phenotypes, including head development, brain development, muscle development, and cellular response to nitrogen (FDR = 0.05; Fig. 4A, Table S5).

**Fig. 4.**
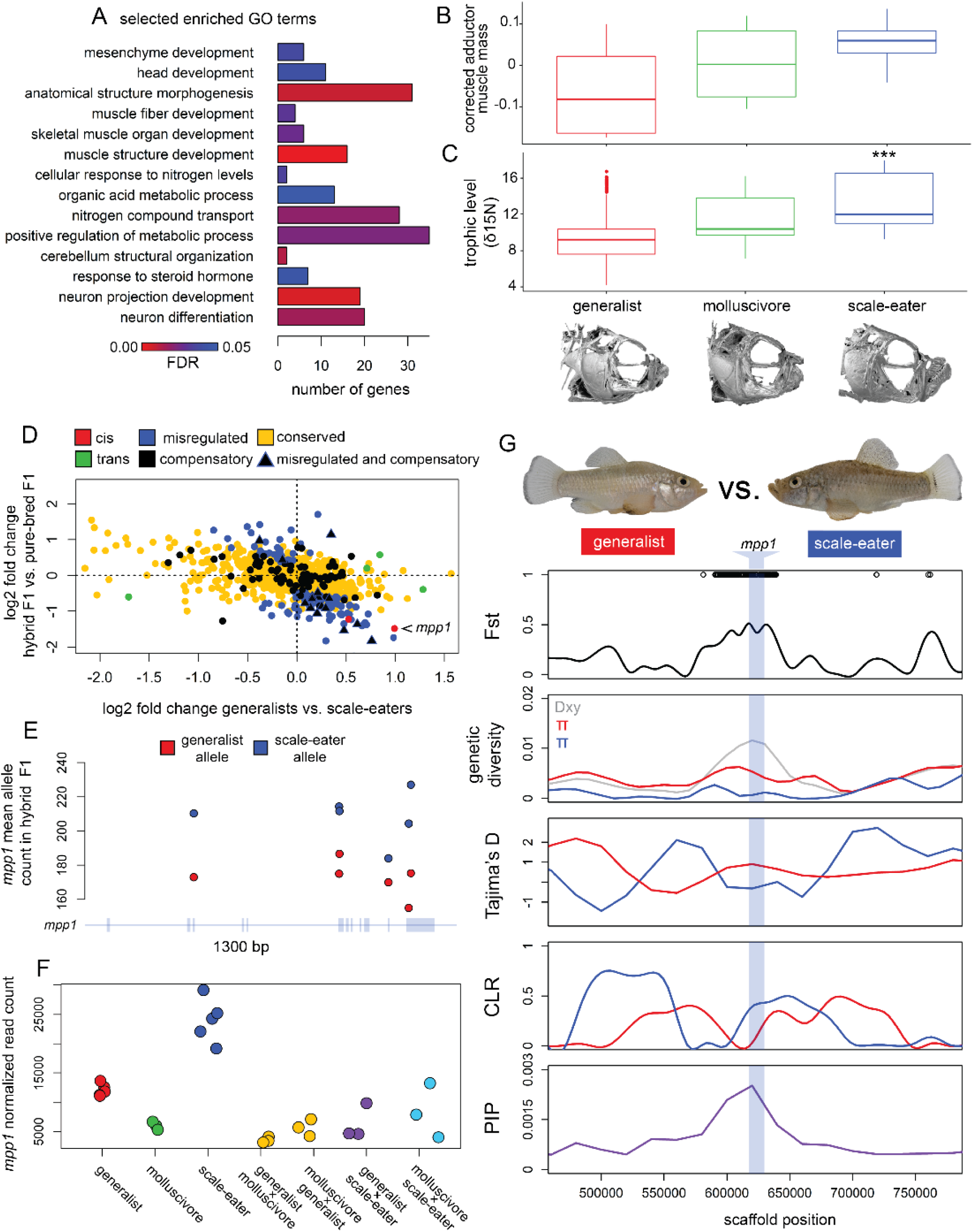
Ecological divergence causes hybrid gene misregulation. A) 14 selected gene ontology (GO) terms relevant to trophic specialization were significantly enriched for the set of 125 genes in highly differentiated genomic regions that showed differential expression between species and misregulation in F1 hybrids. Consistent with muscle development and nitrogen metabolism enrichment, B) adductor mandibulae muscle mass tends to be larger in specialists and C) stable nitrogen isotope ratios (δ15N) are significantly higher in scale-eaters, indicating that they occupy a higher trophic level (Tukey post-hoc test: *P* < 0.001***). D) The gene *mpp1* is controlled by *cis*-regulatory divergence as shown by E) allele specific expression in F1 hybrids and F) differential expression between Crescent Pond generalists vs. scale-eaters and misregulation in their F1 hybrids. G) The gene *mpp1* (light blue band) is near 170 SNPs fixed between Crescent Pond generalists vs. scale-eaters (black points), shows high absolute divergence between species (*D_xy_*), low within-species diversity (π), signatures of a hard selective sweep (Tajima’s D and SweeD composite likelihood ratio (CLR)), and is significantly associated with oral jaw length (PIP; GEMMA genome-wide association mapping).

26 (20.8%) of these ecological DMI candidate genes showed strong evidence of a hard selective sweep in specialists (negative Tajima’s D < genome-wide 10^th^ percentile; range: −1.62 – −0.77; SweeD composite likelihood ratio > 90^th^ percentile by scaffold; Table S6 and S7) and 16 of these showed at least a two-fold expression difference in F1 hybrids compared to purebred F1. Several ecological DMI candidate genes have known functions that are compelling targets for divergent ecological selection. For example, the autophagy-related gene *map1lc3c* has been shown to influence growth when cells are nitrogen deprived (37, 38). Given that specialists occupy higher trophic levels than generalists, as shown by stable isotope ratios (δ15N; Fig. 5B), expression changes in this gene may be important adaptations to nitrogen-rich diets. Similarly, expression changes in the ten genes annotated for effects on brain development may influence divergent behavioral adaptations associated with trophic specialists, including significantly increased aggression (39) and female mate preferences (40).

Using a genome-wide association mapping method that accounts for genetic structure among populations (41), we found that nine of the 125 genes in differentiated regions were significantly associated with oral jaw size – the most rapidly diversifying skeletal trait in this radiation (GEMMA PIP > 99^th^ percentile; Table S8; Fig. S5). For example, we found that *mpp1* was near 170 SNPs fixed between Crescent Pond generalists and scale-eaters, showed evidence of a hard selective sweep in both populations, and was differentially expressed due to *cis-* regulatory mechanisms (Fig. 4F-I). F1 hybrids showed a 3-fold decrease in expression of *mpp1* (*P* = 0.001; Fig. 4F). Knockouts of this gene were recently shown to cause severe craniofacial defects in humans and mice (42). The other eight genes significantly associated with jaw size have not been previously shown to influence cranial phenotypes, but some have known functions in cell types relevant to craniofacial development (Table S8). For example, the gene *sema6c*, which shows strong signs of selection in both scale-eaters and molluscivores (Fig. S6), is known to be expressed at neuromuscular junctions and is important for neuron growth and development within skeletal muscle (43). Expression changes in this gene may influence the development of jaw closing muscles (adductor mandibulae), which tend to be larger in specialists relative to generalists (Fig. 5B). Overall, we found candidate regulatory variants under selection that likely contribute to hybrid gene misregulation and demonstrate that genes near these variants are strikingly enriched for developmental functions related to divergent adaptive traits.

## Discussion

By combining whole genome sequencing with transcriptomic analyses of developing tissues in recently diverged trophic specialists and their F1 hybrids, we provide a genome-wide view of how ecological selection can directly result in genetic incompatibilities causing gene misregulation in hybrids, even in sympatry. Our results are consistent with negative epistatic interactions between alleles from different parental genomes affecting 750 genes (3% of the transcriptome) that show differential expression between species and misregulation in F1 hybrids. 125 of these genes were in highly differentiated regions of the genome containing SNPs fixed between specialists which were enriched for developmental processes relevant to trophic specialization, suggesting that misregulation of these candidate genes in F1 and later generations of hybrids may disrupt the function of adaptive traits and contribute to reproductive isolation between these nascent species.

The negative fitness consequences associated with hybrid gene misregulation are well documented in many systems (14–16, 44, 45), but most of this research has focused on genes associated with sterility and inviability between highly divergent species (but see (23)). It is clear that these strong intrinsic postzygotic isolating barriers evolve more slowly than premating barriers (11, 46, 47); however, hybrid gene misregulation may also have non-lethal effects on fitness and performance that could evolve before or alongside premating isolating mechanisms. Additionally, if genes that are differentially expressed between species in developing tissues are important for adaptive trait divergence, then misregulation of those genes could contribute to abnormal phenotypes that are ecologically maladaptive (18, 23, 48). We previously found extensive gene misregulation specific to craniofacial tissues, which were dissected from generalist × molluscivore F1 hybrids at an early developmental stage (49). Furthermore, F2 and later generation hybrids showing more transgressive phenotypes exhibited the lowest survival and growth rate in field enclosures across multiple lakes and multiple independent field experiments on San Salvador Island (27, 50). In the lab, generalist × scale-eater F1 hybrids exhibited non-additive and impaired feeding performance on scales (28). Overall, these independent lines of evidence suggest that hybrids among San Salvador Island species suffer reduced performance and survival in both laboratory and field environments, which may result from misregulation of genes that are necessary for the normal development of their adaptive traits.

If divergent ecological selection on adaptive traits also causes gene misregulation and subsequently reduced performance and survival of hybrids in the wild, then these ecological DMIs may promote rapid speciation, analogous to the mechanism of magic traits (51). For example, whereas magic traits contribute to RI through assortative mating as a byproduct of divergent ecological selection, these ecological DMIs contribute to RI through gene misregulation and reduced hybrid fitness (18). Thus, our results support a mechanism for divergent ecological selection to generate RI as a byproduct since many adaptive traits are expected to evolve by divergent gene regulation that may come into conflict in a hybrid genetic background (9, 18).

Mathematical models and simulations suggest that genetic incompatibilities evolve most rapidly under directional selection (52, 53), and evolve more slowly under stabilizing selection when compensatory *cis* and *trans* variants have opposing effects on expression levels (52). We see evidence for both types of selection driving misregulation. 22.3% of all misregulated genes showed expression patterns consistent with compensatory regulation, a signature of stabilizing selection (Table S2). However, 26 ecological DMI candidate genes in highly differentiated genomic regions showed strong evidence of hard selective sweeps due to directional selection (Table S6), and more genes may have experienced soft sweeps that were not detected by our methods. Although scale-eaters from Crescent Pond and Osprey Lake form a monophyletic group (Fig. S1), we found little overlap in misregulated genes between lakes (Fig. 2). This may result from selection on Caribbean-wide standing genetic variation that has similar effects on expression, as we showed previously (33), and could reflect polymorphic incompatibilities segregating within species (19). We also see distinct intraspecific differences between lake populations of trophic specialists in pigmentation, maxillary protrusion, and other traits (54), consistent with divergent regulatory variation underlying these adaptive phenotypes.

Identifying genetic variation that contributes to adaptive variation and studying its effect on reproductive isolation is important to understand the sequence of molecular changes leading to ecological speciation. We show that ecologically relevant genes near differentiated genetic regions between sympatric species are under selection and misregulated in F1 hybrids. Overall, our results are consistent with previous observations that hybrid incompatibility alleles are often segregating within populations (17, 19, 55, 56) and that hundreds of genetic incompatibilities can contribute to reproductive isolation between species at the earliest stages of divergence (21). We extend this emerging consensus by showing that gene misregulation can result as a byproduct of divergent ecological selection on a wide range of adaptive traits.

## Methods

### Study system and sample collection

We collected 51 wild-caught individuals from nine isolated hypersaline lakes on San Salvador Island, Bahamas, plus outgroup populations across the Caribbean (see supplemental methods). Our total mRNA transcriptomic dataset consisted of 124 *Cyprinodon* exomes from lab-reared embryos collected between 2017 and 2018. We collected fishes for breeding from two hypersaline lakes on San Salvador Island, Bahamas (Osprey Lake and Crescent Pond); Lake Cunningham, New Providence Island, Bahamas; and Fort Fisher, North Carolina, United States.

We performed 11 separate crosses falling into three categories. 1) For purebred crosses, we collected F1 embryos from breeding tanks containing multiple breeding pairs from a single location. 2) For San Salvador species crosses, we crossed a single individual of one species with a single individual of another species from the same lake for all combinations of the three San Salvador species. In order to control for maternal effects on gene expression inheritance, we collected samples from reciprocal crosses for three of the San Salvador species crosses. 3) For outgroup generalist crosses, we crossed a Crescent Pond generalist male with a Lake Cunningham female and a North Carolina female (Table S9).

### Sequencing and variant discovery

Genomic resequencing libraries were prepared using TruSeq library preparation kits and sequenced on Illumina 150PE Hiseq4000. We mapped a total of 1,953,034,511 adaptor-trimmed reads to the *Cyprinodon* reference genome (57) with the Burrows-Wheeler Alignment Tool (58). We extracted RNA from a total of 348 individuals across two early developmental stages (2 days post fertilization (dpf) and 8 dpf) using RNeasy Mini Kits (Qiagen, Inc.). For 2 dpf libraries, we pooled 5 embryos together and pulverized them in a 1.5 ml Eppendorf tube. We used the same extraction method for samples collected at 8 dpf but did not pool larvae. Libraries were prepared using TruSeq stranded mRNA kits and sequenced on 3 lanes of Illumina 150 PE Hiseq4000 at the Vincent J. Coates Genomic Sequencing Center. We mapped 1,638,067,612 adaptor-trimmed reads to the reference genome using the RNAseq aligner STAR with default parameters (59). We did not find a difference between species or outgroup populations for standard quality control measures, (Fig. S7; ANOVA, *P* > 0.1), except for a marginal difference in transcript integrity numbers (Fig. S8; ANOVA, *P* = 0.041) driven by slightly higher transcript quality in North Carolina generalist samples relative to other samples (Tukey post-hoc test: *P* = 0.043). We found no significant differences among San Salvador Island generalists, molluscivores, scale-eaters, and outgroups in the proportion of reads that mapped to annotated features of the *Cyprinodon* reference genome (Fig. S9; ANOVA, *P* = 0.17).

We used the Genome Analysis Toolkit (60) to call and refine SNP variants across 58 *Cyprinodon* genomes and across 124 *Cyprinodon* exomes. We filtered both SNP datasets to include individuals with a genotyping rate above 90% and SNPs with minor allele frequencies higher than 5%. Our final filtered genomic SNP dataset included 13,838,603 variants with a mean sequencing coverage of 8.2× per individual. We further refined our transcriptomic SNP dataset using the allele-specific software WASP (v. 0.3.3) to correct for potential mapping biases that would influence tests of allele-specific expression (61, 62). We re-called SNPs using unbiased BAMs determined by WASP for a final transcriptomic SNP dataset that included 413,055 variants with a mean coverage of 1,060× across features per individual.

### Phylogenetic analyses

In order to determine the relationship between expression divergence, F1 hybrid misregulation, and phylogenetic distance, we estimated a maximum likelihood tree using RAxML (63). We excluded all missing sites and sites with more than one alternate allele from our genomic SNP dataset, leaving 1,737,591 variants across 58 individuals for analyses. We performed ten separate searches with different random starting trees under the GTRGAMMA model. Node support was estimated from 1,000 bootstrap samples. We fit phylogenetic generalized least-squares (PGLS) models in R with the packages ape (64) and nlme to assess whether gene expression patterns were associated with geographic distance among populations after accounting for phylogenetic relatedness among populations and species. We excluded Osprey Lake populations from these analyses because outgroups were only crossed with Crescent Pond generalists.

### Population genomics and genome-wide association mapping

If alleles causing gene expression divergence between species affect the development of adaptive traits, and also cause gene misregulation in hybrids resulting in low fitness, we predicted that genomic regions near these genes would be strongly differentiated between species, associated with divergent ecological traits, and show signatures of positive selection. We measured relative genetic differentiation (*F_st_*), within population diversity (π), and between population divergence (*D_xy_*) across 58 *Cyprinodon* individuals using 13.8 million SNPs (Table S1 and S7). We identified 20 kb genomic windows significantly associated with variation in oral jaw size across all populations in our dataset (Table S8; Fig. S5). We measured upper jaw lengths and standard length for all individuals in our genomic dataset using digital calipers, fit a log-transformed jaw length by log-transformed standard length linear regression to correct for body size, and used the residuals for genome-wide association mapping with the software GEMMA (41). This program accounts for population structure by incorporating a genetic relatedness matrix into a Bayesian sparse linear mixed model which calculates a posterior inclusion probability (PIP) indicating the proportion of Markov Chain Monte Carlo iterations in which a SNP was estimated to have a non-zero effect on phenotypic variation. We used Tajima’s D statistic and the software SweeD (65) to identify shifts in the site frequency spectrum characteristic of hard selective sweeps. We performed gene ontology enrichment analyses for candidate gene sets using ShinyGo (66).

### Hybrid misregulation and inheritance of gene expression patterns

We aggregated read counts with featureCounts (67) at the transcript isoform level (36,511 isoforms corresponding to 24,952 protein coding genes). Significant differential expression between groups was determined with DESeq2 (68) using Wald tests comparing normalized posterior log fold change estimates and correcting for multiple testing using the Benjamini–Hochberg procedure with a false discovery rate of 0.05 (69). We compared expression in F1 hybrids to expression in F1 purebred offspring to determine whether genes showed additive, dominant, or transgressive patterns of inheritance in hybrids. To categorize hybrid inheritance for F1 offspring generated from a cross between a female from population A and a male from population B (F1_(A×B)_), we conducted four pairwise differential expression tests with DESeq2: 1) F1 _(A)_ vs. F1 _(B)_ 2) F1 _(A)_ vs. F1 _(A×B)_ 3) F1 _(B)_ vs. F1 _(A×B)_ 4) F1 _(A)_ + F1 _(B)_ vs. F1 _(A×B)_. Hybrid inheritance was considered additive if hybrid gene expression was intermediate between parental populations and significantly different between parental populations. Inheritance was dominant if hybrid expression was significantly different from one parental population but not the other. Genes showing misregulation in hybrids showed transgressive inheritance, meaning that hybrid gene expression was significantly higher (overdominant) or lower (underdominant) than both parental species (Fig. S10-12).

### Parallel changes in gene expression in specialists

Parallel evolution of gene expression is often associated with convergent niche specialization, but parallel changes in expression may also underlie divergent specialization (33). We looked at the intersection of genes differentially expressed between generalists versus molluscivores and generalists versus scale-eaters to determine whether both specialists showed parallel changes in expression relative to generalists. We asked whether significant parallelism at the level of gene expression in specialists was mirrored by parallel regulatory mechanisms. We predicted that genes showing parallel changes in specialists would show conserved expression levels in specialist hybrids if they were controlled by the same (or compatible) regulatory mechanisms, but would be misregulated in specialist hybrids if expression was controlled by incompatible regulatory mechanisms. We identified genes showing conserved levels of expression in specialist hybrids (no significant difference in expression between F1 purebreds and F1 hybrids) and genes showing misregulation in specialist hybrids. We also identified genes showing misregulation in specialists relative to all other samples in our dataset across the Caribbean.

### Allele specific expression

Our genomic dataset included every parent used to generate F1 hybrids between populations (*n* = 15). To categorize mechanisms of regulatory divergence between two populations, we used custom R and python scripts (github.com/joemcgirr/fishfASE) to identify SNPs that were alternatively homozygous in breeding pairs and heterozygous in their F1 offspring. We counted reads across heterozygous sites using ASEReadCounter and matched read counts to maternal and paternal alleles. We identified significant ASE using a beta-binomial test comparing the maternal and paternal counts at each gene transcript with the R package MBASED (70). A transcript was considered to show ASE if it showed significant ASE in all F1 hybrid samples generated from the same breeding pair and did not show significant ASE in purebred F1 offspring from the same parental populations.

## Acknowledgements

This study was funded by the University of North Carolina at Chapel Hill, the Miller Institute for Basic Research in the Sciences, NSF CAREER Award 1749764, and NIH/NIDCR R01 DE027052 to CHM. Travel was supported by an SSE Rosemary Grant Award to JAM. We thank Aaron Comeault, Chris Willett, and Jennifer Coughlan for helpful comments on the manuscript; Daniel Matute, Emilie Richards, Michelle St. John, Bryan Reatini, and Sara Suzuki for valuable discussion; The Vincent J. Coates Genomics Sequencing Laboratory at the University of California, Berkeley for performing RNA library prep and Illumina sequencing; the Gerace Research Centre for logistics; and the Bahamian government BEST Commission for permission to conduct this research.

## Data Availability

All transcriptomic raw sequence reads are available as zipped fastq files on the NCBI BioProject database. Accession: PRJNA391309. Title: Craniofacial divergence in Caribbean Pupfishes. All R and Python scripts used for pipelines are available on Git (github.com/joemcgirr/fishfASE).

## Supplemental Methods

### Study system and sample collection

We collected 51 wild-caught individuals from nine isolated hypersaline lakes on San Salvador Island, Bahamas (Great Lake, Stout’s Lake, Oyster Lake, Little Lake, Crescent Pond, Moon Rock, Mermaid’s Pond, Osprey Lake, Pigeon Creek) between 2011 and 2018 using seine-nets and hand nets. 18 scale-eaters (*Cyprinodon desquamator*) were sampled from six lake populations; 15 molluscivores (*C. brontotheroides*) were sampled from four populations; and 18 generalists (*C. variegatus*) were sampled from nine populations. The genomic dataset also included two *C. laciniatus* from Lake Cunningham, New Providence Island, Bahamas, one *C. bondi* from Etang Saumautre lake in the Dominican Republic, one *C. variegatus* from Fort Fisher, North Carolina, one *C. diabolis* from Devils Hole, Nevada, and captive-bred individuals of *C. simus* and *C. maya* from Laguna Chicancanab, Quintana Roo, Mexico. Sampling is further described in (1, 2). Fish were euthanized in an overdose of buffered MS-222 (Finquel, Inc.) following approved protocols from the University of California, Davis Institutional Animal Care and Use Committee (#17455), the University of California, Berkeley Animal Care and Use Committee (AUP-2015-01-7053), and the University of North Carolina Institutional Animal Care and Use Committee (18-061.0). Samples were stored in 95-100% ethanol.

Our total mRNA transcriptomic dataset consisted of 124 *Cyprinodon* exomes from embryos collected between 2017 and 2018. We collected fishes for breeding from two hypersaline lakes on San Salvador Island, Bahamas (Osprey Lake, and Crescent Pond), Lake Cunningham, New Providence Island, Bahamas, and Fort Fisher, North Carolina, United States.. Wild-caught parents were reared in breeding tanks at 25–27°C, 10–15 ppt salinity, pH 8.3, and fed a mix of commercial pellet foods and frozen foods. All purebred F1 offspring were collected from breeding tanks containing multiple F0 breeding pairs. All F1 offspring from crosses between species and populations were collected from individual F0 breeding pairs that were subsequently sequenced in our genomic dataset.

Methods for collecting and raising embryos were similar to previously outlined methods (3, 4). All F1 embryos were collected from breeding mops within one hour of spawning and transferred to petri dishes incubated at 27°C. Embryo water was treated with Fungus Cure (API Inc.) and changed every 48 hours. Embryos were inspected for viability and sampled either 47-49 hours post fertilization (hereafter 2 days post fertilization (2 dpf)) or 190-194 hours (eight days) post fertilization (hereafter 8 dpf). These early developmental stages are described as stage 23 (2 dpf) and 34 (8 dpf) in a recent embryonic staging series of *C. variegatus* (5). The 2 dpf stage is comparable to the Early Pharyngula Period of zebrafish, when multipotent neural crest cells have begun migrating to pharyngeal arches that will form the oral jaws and most other craniofacial structures (6–8). Embryos usually hatch six to ten days post fertilization, with similar variation in hatch times among species (3, 7). While some cranial elements are ossified prior to hatching, the skull is largely cartilaginous at 8 dpf (5). Embryos from each stage were euthanized in an overdose of buffered MS-222 and immediately preserved in RNA later (Ambion, Inc.) for 24 hours at 4°C and then - 20°C for up to 9 months following manufacturer’s instructions.

### Hybrid cross design

All parents used to generate F1 hybrids were collected from four locations: 1) Crescent Pond, San Salvador, 2) Osprey Lake, San Salvador, 3) Lake Cunningham, New Providence Island, or 4) Fort Fisher, North Carolina. In order to understand how varying levels of genetic divergence and ecological divergence between parents affected gene expression patterns in F1 offspring, we performed 11 separate crosses falling into three categories. 1) For purebred crosses, we collected F1 embryos from breeding tanks containing multiple breeding pairs from a single location. 2) For San Salvador species crosses, we crossed a single individual of one species with a single individual of another species from the same lake for all combinations of the three San Salvador species. In order to control for maternal effects on gene expression inheritance, we collected samples from reciprocal crosses for three San Salvador species crosses. 3) For outgroup generalist crosses, we bred a Crescent Pond generalist male with a Lake Cunningham female and a North Carolina female (Table S9).

### Genomic sequencing and alignment

All DNA samples were extracted from muscle tissue or caudal fin clips using DNeasy Blood and Tissue kits (Qiagen, Inc.) and quantified on a Qubit 3.0 fluorometer (Thermofisher Scientific, Inc.). Sequencing methods for 43 of the 58 individuals in our genomic dataset were previously described (1, 2). Briefly, libraries were prepared using Illumina TruSeq DNA PCR-Free kits at the Vincent J. Coates Genomic Sequencing Center (QB3, Berkeley, CA) and samples were pooled on four lanes of Illumina 150PE Hiseq4000. We added 15 new individuals to this dataset that were crossed to generate F1 hybrids. These libraries were prepared at the same facility using TruSeq kits on the automated Apollo 324 system (WaferGen BioSystems, Inc.). Samples were fragmented using Covaris sonication, barcoded with Illumina indices, quality checked using a Fragment Analyzer (Advanced Analytical Technologies, Inc.), and sequenced on one lane of Illumina 150PE Hiseq4000 in June 2018.

We filtered raw reads using Trim Galore (v. 4.4, Babraham Bioinformatics) to remove Illumina adaptors and low-quality reads (mean Phred score < 20) and mapped 1,953,034,511 reads to the *Cyprinodon* reference genome (NCBI, *Cyprinodon variegatus* annotation release 100; total sequence length = 1,035,184,475; number of scaffolds = 9,259; scaffold N50 = 835,301; contig N50 = 20,803; (7)) with the Burrows-Wheeler Alignment Tool (bwa mem; (9) (v. 0.7.12)). The Picard software package (v. 2.0.1) and Samtools (v. 1.9) were used to remove duplicate reads (MarkDuplicates) and create indexes. We assessed mapping and read quality using MultiQC (10).

### Transcriptomic sequencing and alignment

We extracted RNA from a total of 348 individuals (whole-embryos and whole-larvae) using RNeasy Mini Kits (Qiagen catalog #74104). For samples collected at 2 dpf, we pooled 5 embryos together and pulverized them in a 1.5 ml Eppendorf tube using a plastic pestle washed with RNase Away (Molecular BioProducts). We used the same extraction method for samples collected at 8 dpf but did not pool larvae and prepared a library for each individual separately. Total mRNA sequencing libraries for the resulting 125 samples were prepared at the Vincent J. Coates Genomic Sequencing Center (QB3, Berkeley, CA) using the Illumina stranded Truseq RNA kit (Illumina RS-122-2001). Sequencing was performed on Illumina Hiseq4000 150PE. 72 and 53 total mRNA libraries were each pooled across three lanes and sequenced in May 2018 and July 2018, respectively.

We filtered raw reads using Trim Galore (v. 4.4, Babraham Bioinformatics) to remove Illumina adaptors and low-quality reads (mean Phred score < 20) and mapped 1,638,067,612 filtered reads to the *Cyprinodon* reference genome (NCBI, *Cyprinodon variegatus* annotation release 100; 1.035 Gb; scaffold N50 = 835,301; (7)) using the RNA-seq aligner STAR with default parameters (v. 2.5 (11)). We assessed mapping and read quality using MultiQC (10). We quantified the number of duplicate reads produced during sequence amplification and GC content of transcripts for each sample using RSeQC (12). We also used RSeQC to estimate transcript integrity numbers (TINs) which is a measure of potential *in vitro* RNA degradation within a sample. TIN is calculated by directly analyzing the uniformity of read coverage across a transcript and is a more reliable measure of degradation compared to RNA integrity number (RIN) which uses ribosomal RNA as a proxy for overall RNA integrity (12, 13). We performed one-way ANOVA to determine whether the GC content of reads, read depth across features, total normalized counts, or TINs differed between samples grouped by species and population. We did not find a difference between species or generalist populations for any quality control measure (Fig. S7; ANOVA, *P* > 0.1), except for a marginal difference in TIN (Fig. S8; ANOVA, *P* = 0.041) driven by slightly higher transcript quality in North Carolina samples (Tukey multiple comparisons of means; *P* = 0.043). We found no significant differences among San Salvador Island generalists, molluscivores, scale-eaters, and outgroup generalists in the proportion of reads that map to annotated features of the *Cyprinodon* reference genome (Fig. S9; ANOVA, *P* = 0.17). We did find that more reads mapped to features in 2 dpf samples than 8 dpf samples (Fig. S13; Student’s *t*-test, *P* < 2.2 × 10^-16^).

### Variant discovery and population genetic analyses

We followed the best practices guide recommended by the Genome Analysis Toolkit (v. 3.5 (14)) in order to call and refine SNP variants across 58 *Cyprinodon* genomes and across 124 *Cyprinodon* exomes using the Haplotype Caller function. For both datasets, we used conservative hard filtering criteria to call SNPs (14, 15): Phred-scaled variant confidence divided by the depth of nonreference samples > 2.0, Phred-scaled *P*-value using Fisher’s exact test to detect strand bias > 60, Mann–Whitney rank-sum test for mapping qualities (z > 12.5), Mann– Whitney rank-sum test for distance from the end of a read for those with the alternate allele (z > 8.0). We filtered both SNP datasets to include individuals with a genotyping rate above 90% and SNPs with minor allele frequencies higher than 5%. Our final filtered genomic SNP dataset included 13,838,603 variants with a mean sequencing coverage of 8.2× per individual.

We further refined our transcriptomic SNP dataset using the allele-specific software WASP (v. 0.3.3) to correct for potential mapping biases that would influence tests of allele-specific expression (ASE; (16, 17)). While we showed that mapping bias does not significantly affect the proportion of reads mapped to features between species (Fig. S9), even a small number of biased sites would likely account for the majority of significant ASE at an exome-wide scale. WASP identified reads that overlapped sites in our original transcriptomic SNP dataset and re-mapped those reads after swapping the genotype for the alternate allele. Reads that failed to map to exactly the same location were discarded. We re-mapped unbiased reads using methods outlined above to create our final BAM files that were used for all downstream analyses. We re-called SNPs using unbiased BAMs for a final transcriptomic SNP dataset that included 413,055 variants with a mean coverage of 1,060× across gene features per individual.

We analyzed genomic SNPs to measure within-population diversity (π), between-population diversity (*D_xy_*), relative genetic diversity (*F_st_*), and Tajima’s D. We measured π, *D_xy_*, and *F_st_* in 20 kb windows using the python script popGenWindows.py created by Simon Martin (available on https://github.com/simonhmartin/genomics_general; (18)). 13.8 million SNP variants genotyped by whole genome resequencing of 58 *Cyprinodon* individuals revealed more population structure between allopatric generalists than between generalists and specialists on San Salvador (genome-wide mean *F_st_* between San Salvador generalists: vs. North Carolina = 0.217; vs. New Providence = 0.155; vs. scale-eaters = 0.106; vs. molluscivores = 0.056). We found consistent relationships across a maximum likelihood phylogeny calculated with RAxML, with longer branch lengths separating allopatric populations (Fig. 1, S1).

We calculated Tajima’s D in 20 kb windows and per site *F_st_* for each species and lake population genomic using VCFtools (v. 1.15). We chose to analyze 20 kb windows given previous estimates of pairwise linkage disequilibrium (measured as *r*^2^) showing that linkage dropped to background levels between SNPs separated by >20 kb (*r*^2^<0.1; (1)). Tajima’s D statistic compares observed nucleotide diversity to diversity under a null model assuming genetic drift, where negative values indicate a reduction in diversity across segregating sites that may be due to positive selection (19). We also looked for evidence of hard selective sweeps using the SweepFinder method first developed by Nielsen et al. (2005) and implemented in the software package SweeD (20, 21). SweeD separates scaffolds into 1000 windows of equal size and calculates a composite likelihood ratio (CLR) from a comparison of two contrasting models for each window. The first assumes a window has undergone a recent selective sweep, whereas the second assumes a null model where the site frequency spectrum of the window does not differ from that of the entire scaffold. Windows with a high CLR suggest a history of selective sweeps because the site frequency spectrum is shifted toward low-frequency and high-frequency derived variants (20, 21).

We used ancestral population sizes (previously determined by the Multiple Sequentially Markovian Coalescent approach (1, 22) to estimate the expected neutral SFS with SweeD, accounting for historical demographic effects on the contemporary shape of the SFS. SweeD identifies regions of a scaffold showing signs of a hard sweep relative to the rest of that scaffold. Thus, we normalized CLR values to be between zero and one to compare the strength of selection across scaffolds. We defined regions showing strong signs of a hard selective sweep as windows that showed CLRs above the 90^th^ percentile for a scaffold (normalized CLR > 0.9) and a negative value of Tajima’s D less than the genome-wide 10^th^ percentile (range = −1.62 – −0.77 (see Table S7 for all population thresholds)). We also visually inspected regions near candidate incompatibility genes to identify CLRs and Tajima’s D estimates indicating moderate signs of selection.

### Read count abundance and differential expression analyses

We used the featureCounts function of the Rsubread package (23) requiring paired-end and reverse stranded options to generate read counts across 36,511 previously annotated features for the *Cyprinodon* reference genome (7). We aggregated read counts at the transcript isoform level (36,511 isoforms correspond to 24,952 protein coding genes).

We used DESeq2 (v. 3.5 (24)) to normalize raw read counts and perform principal component analyses. DESeq2 normalizes read counts by calculating a geometric mean of counts for each gene across samples, dividing individual gene counts by this mean, and then using the median of these ratios as a size factor for each sample. These sample-specific size factors account for differences in library size and sequencing depth among samples. Gene features showing less than 10 normalized counts in every sample were discarded from analyses. We constructed a DESeqDataSet object in R using a multi-factor design that accounted for variance in F1 read counts influenced by parental population origin and sequencing date (design = ∼sequencing_date + parental_breeding_pair_populations). Next, we used a variance stabilizing transformation on normalized counts and performed a principal component analysis to visualize the major axes of variation in 2 dpf and 8 dpf samples (Fig. S15). We removed one 8 dpf outlier so that the final count matrix used for differential expression analyses included 124 samples (2 dpf = 68, 8 dpf = 56).

DESeq2 fits negative binomial generalized linear models for each gene across samples to test the null hypothesis that the fold change in gene expression between two groups is zero. The program uses an empirical Bayes shrinkage method to determine gene dispersion parameters, which model within-group variability in gene expression and logarithmic fold changes in gene expression. Significant differential expression between groups was determined with Wald tests by comparing normalized posterior log fold change estimates and correcting for multiple testing using the Benjamini–Hochberg procedure with a false discovery rate of 0.05 (Benjamini and Hochberg 1995). We contrasted gene expression in pairwise comparisons between populations grouped by developmental stage. To determine within population levels of expression divergence (Fig. 1B-E), we down-sampled each population to perform every pairwise comparison between samples using the highest sample size possible between groups and calculated the mean number of genes differentially expressed across comparisons.

### Hybrid misregulation and inheritance of gene expression patterns

We generated F1 hybrid offspring from crosses between populations and generated purebred F1 offspring from crosses within populations. We compared expression in hybrids to expression in purebred offspring to determine whether genes showed additive, dominant, or transgressive patterns of inheritance in hybrids. To categorize hybrid inheritance for F1 offspring generated from a cross between a female from population A and a male from population B (F1_(A×B)_), we conducted four pairwise differential expression tests with DESeq2:

1. F1 _(A)_ vs. F1 _(B)_
2. F1 _(A)_ vs. F1 _(A×B)_
3. F1 _(B)_ vs. F1 _(A×B)_
4. F1 _(A)_ + F1 _(B)_ vs. F1 _(A×B)_

Hybrid inheritance was considered additive if hybrid gene expression was intermediate between parental populations and significantly different between parental populations. Inheritance was dominant if hybrid expression was significantly different from one parental population but not the other. Genes showing misregulation in hybrids showed transgressive inheritance, meaning hybrid gene expression was significantly higher (overdominant) or lower (underdominant) than both parental species (Fig. S10-12). All comparisons were conducted between groups sampled at the same developmental stage (2 dpf or 8 dpf).

### Parallel changes in gene expression in specialists

Parallel evolution of gene expression is often associated with convergent niche specialization, but parallel changes in expression may also underlie divergent specialization (4). We looked at the intersection of genes differentially expressed between generalists versus molluscivores and generalists versus scale-eaters to determine whether specialists showed parallel changes in expression relative to generalists. We compared expression between generalists and each specialist grouping samples by lake population and developmental stage.

We also examined the direction of expression divergence for each gene to evaluate the significance of parallel expression evolution (Fig 3E). Specifically, we wanted to know whether the fold change in expression for genes tended to show the same sign in both specialists relative to generalists (either up-regulated in both specialists relative to generalists or down-regulated in both specialists). Under a neutral model of gene expression evolution, half of the genes differentially expressed between generalists versus molluscivores and generalists versus scale-eaters would show fold changes in the same direction and half would show fold changes in opposite directions (Fig. 3E). Remarkably, 1,206 (96.6%) of the genes showing expression divergence between generalists versus molluscivores and generalists versus scale-eaters showed the same direction of expression divergence in specialists. These results provide robust evidence for parallel changes in expression underlying divergent trophic adaptation and support previous findings based on a smaller sample size (3).

We wanted to determine whether significant parallelism at the level of gene expression in specialists was mirrored by parallel regulatory mechanisms. We predicted that genes showing parallel changes in specialists would show conserved expression levels in specialist hybrids if they were controlled by the same (or compatible) regulatory mechanisms, but would be misregulated in specialist hybrids if expression was controlled by different and incompatible regulatory mechanisms. We identified genes showing conserved levels of expression in specialist hybrids (no significant difference in expression between purebred specialist F1s and specialist hybrid F1s) and genes showing misregulation in specialist hybrids. We also identified genes showing extreme Caribbean-wide misregulation in specialists. These genes were differentially expressed in specialist hybrids relative to all other samples in our dataset from across the Caribbean (North Carolina to New Providence Island, Bahamas).

### Allele specific expression and mechanisms of regulatory divergence

We partitioned hybrid gene expression divergence into patterns that could be attributed to *cis*-regulatory variation in cases where linked genetic variation affected proximal gene expression levels, and *trans*-regulatory variation in cases where genetic variation in unlinked factors bound to *cis*-regulatory elements affected gene expression levels. It is possible to identify mechanisms of gene expression divergence between parental species by bringing *cis* elements from both parents together in the same *trans* environment in F1 hybrids and quantifying allele specific expression (ASE) of parental alleles at heterozygous sites (25, 26). A gene showing ASE in F1 hybrids that is differentially expressed between parental species is expected to result from *cis*-regulatory divergence. *Trans*-regulatory divergence can be determined by comparing the ratio of gene expression in parents with the ratio of allelic expression in F1 hybrids. *Cis* and *trans* regulatory variants often interact to affect expression divergence of the same gene (26–28).

Our genomic variant dataset included every parent used to generate F1 hybrids between populations (*n* = 15). We used the VariantsToTable function of the Genome Analysis Toolkit (14) to output genotypes across 13.8 million variant sites for each parent and overlapped these sites with the 413,055 variant sites identified across F1 transcriptomes (corrected for mapping bias). To categorize mechanisms of regulatory divergence between two populations, we used custom R and python scripts (https://github.com/joemcgirr/fishfASE) to identify SNPs that were alternatively homozygous in breeding pairs and heterozygous in their F1 offspring. We counted reads across heterozygous sites using ASEReadCounter (-minDepth 20 --minMappingQuality 10 --minBaseQuality 20 -drf DuplicateRead) and matched read counts to maternal and paternal alleles. We calculated the significance of ASE per gene transcript. We identified significant ASE using a beta-binomial test comparing the maternal and paternal counts at each transcript with the R package MBASED (29). For each F1 hybrid sample, we performed a 1-sample analysis with MBASED using default parameters run for 1,000,000 simulations to identify transcripts showing significant ASE (*P* < 0.05). Finally, we quantified allele counts across all heterozygous sites for each purebred F1 sample and ran the same analyses in MBASED to identify transcripts showing ASE in parental populations. A transcript was considered to show ASE if it showed significant ASE in all F1 hybrid samples generated from the same breeding pair and did not show significant ASE in purebred F1 offspring generated from the same parental populations.

In order to determine regulatory mechanisms controlling expression divergence between parental species, a transcript had to be included in differential expression analyses and ASE analyses. We were able to classify regulatory categories for more transcripts if breeding pairs were more genetically divergent because we could analyze more heterozygous sites in their hybrids (mean number of informative transcripts across crosses = 1,914; range = 812 – 3,543). For each hybrid sample and each transcript amenable to both types of analyses, we calculated H – the ratio of maternal allele counts compared to the number of paternal allele counts in F1 hybrids, and P – the ratio of normalized read counts in purebred F1 offspring from the maternal population compared to read counts in purebred F1 offspring from the paternal population. We performed a Fisher’s exact test using H and P to determine whether there was a significant *trans-* contribution to expression divergence, testing the null hypothesis that the ratio of read counts in the parental populations was equal to the ratio of parental allele counts in hybrids (26, 28, 30, 31).

We classified expression divergence due to *cis*-regulation if a transcript showed significant ASE, significant differential expression between parental populations of purebred F1 offspring, and no significant *trans*-contribution. We identified expression divergence due to *trans*-regulation if transcripts did not show ASE, were differentially expressed between parental populations of purebred F1 offspring, and showed significant *trans*-contribution. We found compensatory regulatory divergence (*cis-* and *trans*-regulatory factors had opposing effects on expression) in cases where a transcript showed ASE and was not differentially expressed between parental populations of purebred F1 offspring (Fig. S2-S4).

### Phylogenetic analyses

Gene expression evolves under the combined forces of selection and drift, and is expected to diverge linearly with increasing phylogenetic distance between closely related species (32). The magnitude of F1 hybrid misregulation likely also depends on phylogenetic distance between parental species (33). In order to determine the relationship between expression divergence, hybrid misregulation, and phylogenetic distance, we constructed a maximum likelihood tree using RAxML. We excluded all missing sites and sites with more than one alternate allele from our genomic SNP dataset, leaving 1,737,591 variants across 58 individuals for analyses. We performed ten separate searches with different random starting trees under the GTRGAMMA model. Node support was estimated from 1,000 bootstrap samples. We used branch lengths from the best fitting tree as a measure of phylogenetic distance between populations.

We tested whether isolation by distance (kilometers separating populations) was a significant predictor of gene expression divergence between populations. We also tested whether isolation by distance explained patterns of misregulation in hybrids generated by inter-population crosses. Gene expression levels between species cannot be considered to be independent and identically distributed random variables (34). We used phylogenetic generalized least-squares (PGLS) models in R, using the packages ape (35) and nlme to assess whether gene expression patterns were predicted by distance between populations (measured in kilometers) after accounting for phylogenetic relatedness. We excluded Osprey Lake populations from these analyses because outgroup generalist hybrid crosses only involved Crescent Pond generalists. We used lake diameter as the distance between populations for sympatric comparisons.

### Morphometrics

We used digital calipers to measure upper oral jaw length and body length from external landmarks on ethanol-preserved tissue specimens. Upper jaw length was measured from the quadroarticular joint to the tip of the most anterior tooth on the dentigerous arm of the premaxilla. Body length was measured from the midline of the posterior margin of the caudal peduncle to the tip of the lower jaw. We used this measure of body length rather than standard length to account for size variation because the nasal protrusion on some molluscivore samples extended beyond the upper jaw. One scale-eater specimen was removed from the analysis because the caudal region was missing, preventing an accurate measure of body length. All jaw length measurements were log-transformed and regressed against log-transformed body length to remove the effects of size variation among specimens. Size-corrected residuals were used for genome-wide association mapping

### Association mapping

We employed a Bayesian Sparse Linear Mixed Model (BSLMM) implemented in the GEMMA software package ((36) v. 0.94.1) to identify genomic regions associated with variation in upper oral jaw length. We previously used this program to identify candidate genes influencing jaw size (1). Here, we used the same methods adding 15 individuals to our genomic dataset. Briefly, the BSLMM uses Markov Chain Monte Carlo sampling to estimate the proportion of phenotypic variation explained by every SNP included in the analysis (PVE), the proportion of phenotypic variation explained by SNPs of large effect (PGE), which are defined as SNPs with a non-zero effect on the phenotype, and the number of large-effect SNPs needed to explain PGE (nSNPs; Fig. S5). GEMMA also estimates an effect size coefficient (β) and a posterior inclusion probability (PIP) for each SNP. We used PIP (the proportion of iterations in which a SNP is estimated to have a non-zero effect on phenotypic variation (β ≠ 0)) to assess the significance of regions associated with jaw size variation. Because these statistics are difficult to interpret for causal SNPs tightly linked to neutral SNPs, we summed β and PIP parameters across 20-kb windows to avoid dispersion of the posterior probability density across SNPs in linkage disequilibrium (LD). Pairwise LD (*r*^2^) drops to background levels of LD between SNPs separated by more than 20 kb (1). GEMMA controls for background population structure by estimating and incorporating a kinship relatedness matrix as a covariate in the regression model. We performed 10 independent runs of the BSLMM for 57 individuals (following (37)) using a step size of 100 million with a burn-in of 50 million steps. Independent runs were consistent in reporting the strongest associations for the same 20 kb windows. Windows that showed PIP values above the 99th percentile (0.00175) were considered to be strongly associated with oral jaw size variation within Caribbean pupfishes. Our PIP estimates for strongly associated windows suggest that jaw length may be controlled by several loci of moderate effect (see bimodal PGE distribution, Fig. S5B). Indeed, a linkage mapping analysis of phenotypic diversity in an F_2_ intercross between specialists estimated four QTL with moderate effects on oral jaw size explaining up to 15% of the variation (38). Encouragingly, the window that showed the strongest association with jaw size (PIP = 0.1043; Fig. S5) contained a single gene associated with craniofacial deformities in humans (*samd12*; (39)). Additionally, *clk2, gpr119, doc2b, rapgef4*, were also within the top four windows showing the highest PIP values.

### Gene ontology enrichment analyses

We performed a gene ontology (GO) enrichment analysis for the 125 genes in differentiated genomic regions showing differential expression between species and misregulation in hybrids using ShinyGo v.0.51 (40). The RefSeq genome records for the *Cyprinodon* reference genome were annotated by the NCBI Eukaryotic Genome Annotation Pipeline, an automated pipeline that annotates genes, transcripts and proteins. Gene symbols for orthologs identified by this pipleline largely match human gene symbols. Thus, we searched for enrichment across biological process ontologies curated for human gene functions. We also determined whether genes sets showing other interesting patterns of expression were annotated for effects on cranial skeletal system development (GO:1904888).

---

**Table S1.** San Salvador Island population genomic statistics measured across 13.8 million SNPs. Statistics for the top three rows were calculated for all San Salvador individuals of each species (see Fig. S1). The remaining rows are comparisons separated by lake populations used to generate samples for RNAseq (CP = Crescent Pond, OL = Osprey Lake).

| population 1 | n | population 2 | n | mean $D_{xy}$ | $D_{xy}$ 90th percentile | mean $F_{st}$ | # fixed SNPs |
| --- | --- | --- | --- | --- | --- | --- | --- |
| all generalists | 8 | all molluscivores | 10 | 0.0047 | 0.0076 | 0.0564 | 179 |
| all generalists | 8 | all scale-eaters | 9 | 0.0047 | 0.0080 | 0.1065 | 5,331 |
| all molluscivores | 10 | all scale-eaters | 9 | 0.0049 | 0.0085 | 0.1357 | 36,335 |
| CP generalists | 5 | CP molluscivores | 5 | 0.0042 | 0.0075 | 0.0740 | 11,015 |
| CP generalists | 5 | CP scale-eaters | 5 | 0.0046 | 0.0082 | 0.1356 | 109,072 |
| CP molluscivores | 5 | CP scale-eaters | 5 | 0.0048 | 0.0093 | 0.1839 | 559,728 |
| OL generalists | 3 | OL molluscivores | 5 | 0.0049 | 0.0084 | 0.0964 | 47,356 |
| OL generalists | 3 | OL scale-eaters | 4 | 0.0049 | 0.0084 | 0.1130 | 108,813 |
| OL molluscivores | 5 | OL scale-eaters | 4 | 0.0049 | 0.0087 | 0.1347 | 168,192 |
| CP generalists | 5 | OL generalists | 3 | 0.0049 | 0.0082 | 0.0759 | 19,582 |
| CP molluscivores | 5 | OL molluscivores | 5 | 0.0045 | 0.0082 | 0.1169 | 92,317 |
| CP scale-eaters | 5 | OL scale-eaters | 4 | 0.0035 | 0.0073 | 0.0983 | 86,367 |

**Table S2.** Percentage of genes controlled by different regulatory mechanisms for each hybrid cross. Informative genes are those containing heterozygous sites in hybrids that were alternatively homozygous in parents. The final column is the percentage of misregulated genes showing no difference in expression between parental populations and allele-specific expression in F1 hybrids, consistent with compensatory regulatory divergence. NC = North Carolina, NP = New Providence, CP = Crescent Pond, OL = Osprey Lake.

| **mother** | **father** | **stage** | **informative**<br>**genes** | **conserved** | **cis** | **trans** | **compensatory** | **misregulated** | **misregulated**<br>**showing**<br>**compensatory** |
| --- | --- | --- | --- | --- | --- | --- | --- | --- | --- |
| NC generalist | CP generalist | 2dpf | 2182 | 61.18 | 2.66 | 0.37 | 19.98 | 15.81 | 32.75 |
| NP generalist | CP generalist | 2dpf | 2359 | 79.57 | 0.34 | 0.42 | 16.45 | 3.22 | 11.84 |
| CP generalist | CP molluscivore | 2dpf | 2703 | 60.82 | 0.18 | 0.26 | 33.70 | 5.03 | 37.50 |
| CP generalist | CP scale-eater | 2dpf | 1764 | 83.50 | 0.17 | 0.17 | 15.87 | 0.28 | 40.00 |
| CP molluscivore | CP scale-eater | 2dpf | 1645 | 69.79 | 1.64 | 0.55 | 26.38 | 1.64 | 33.33 |
| CP molluscivore | CP generalist | 2dpf | 2193 | 62.79 | 0.14 | 0.05 | 36.07 | 0.96 | 57.14 |
| OL generalist | OL molluscivore | 2dpf | 3114 | 46.66 | 0.03 | 0.06 | 34.20 | 19.04 | 38.95 |
| OL generalist | OL scale-eater | 2dpf | 1934 | 62.77 | 0.05 | 0.52 | 22.75 | 13.91 | 18.96 |
| OL scale-eater | OL molluscivore | 2dpf | 3485 | 74.09 | 1.03 | 0.69 | 23.39 | 0.80 | 21.43 |
| OL molluscivore | OL generalist | 2dpf | 2915 | 59.79 | 0.03 | 0.03 | 37.29 | 2.85 | 38.55 |
| OL scale-eater | OL generalist | 2dpf | 2377 | 57.72 | 0.21 | 0.59 | 29.66 | 11.82 | 31.32 |
| NC generalist | CP generalist | 8dpf | 2995 | 60.13 | 1.40 | 0.07 | 8.41 | 29.98 | 13.47 |
| NP generalist | CP generalist | 8dpf | 1406 | 93.24 | 0.21 | 0.71 | 5.41 | 0.43 | 50.00 |
| CP generalist | CP molluscivore | 8dpf | 819 | 87.06 | 0.12 | 0.12 | 9.77 | 2.93 | 20.83 |
| CP generalist | CP scale-eater | 8dpf | 1147 | 81.17 | 0.26 | 0.44 | 6.63 | 11.51 | 13.64 |
| CP molluscivore | CP scale-eater | 8dpf | 1027 | 78.87 | 1.85 | 2.04 | 4.48 | 12.76 | 8.40 |
| CP molluscivore | CP generalist | 8dpf | 1327 | 88.55 | 0.08 | 0.15 | 10.55 | 0.68 | 33.33 |
| OL generalist | OL molluscivore | 8dpf | 1322 | 75.26 | 0.45 | 0.61 | 7.19 | 16.49 | 9.63 |
| OL generalist | OL scale-eater | 8dpf | 1273 | 85.62 | 0.24 | 1.57 | 3.38 | 9.19 | 13.68 |
| OL scale-eater | OL molluscivore | 8dpf | 984 | 90.24 | 0.81 | 0.61 | 6.20 | 2.13 | 14.29 |
| OL scale-eater | OL generalist | 8dpf | 1087 | 73.60 | 0.18 | 1.10 | 2.21 | 22.91 | 5.62 |

**Table S3.** Number of genes showing differential expression (DE) between species and misregulation in F1 hybrids. Lines separate cross type (top: specialists, middle: generalist and scale-eater, bottom: generalist and molluscivore).

| maternal population | paternal population | informative genes | DE between species | misregulated in F1 | DE and misregulated | stage |
| --- | --- | --- | --- | --- | --- | --- |
| CP molluscivore | CP scale-eater | 11718 | 862 | 88 | 10 | 2dpf |
| OL scale-eater | OL molluscivore | 11820 | 1900 | 150 | 32 | 2dpf |
| CP molluscivore | CP scale-eater | 13013 | 4141 | 1208 | 320 | 8dpf |
| OL scale-eater | OL molluscivore | 13225 | 2020 | 158 | 18 | 8dpf |
| CP generalist | CP scale-eater | 11671 | 335 | 7 | 0 | 2dpf |
| OL generalist | OL scale-eater | 11650 | 1455 | 1453 | 362 | 2dpf |
| CP generalist | CP scale-eater | 13300 | 716 | 1009 | 87 | 8dpf |
| OL generalist | OL scale-eater | 13254 | 3918 | 1088 | 244 | 8dpf |
| OL scale-eater | OL generalist | 11650 | 1455 | 1283 | 38 | 2dpf |
| OL scale-eater | OL generalist | 13254 | 3918 | 2016 | 72 | 8dpf |
| CP generalist | CP molluscivore | 12202 | 606 | 536 | 37 | 2dpf |
| OL generalist | OL molluscivore | 12207 | 97 | 2142 | 4 | 2dpf |
| CP generalist | CP molluscivore | 13594 | 371 | 168 | 13 | 8dpf |
| OL generalist | OL molluscivore | 13697 | 1945 | 1780 | 194 | 8dpf |
| CP molluscivore | CP generalist | 11814 | 606 | 69 | 4 | 2dpf |
| OL molluscivore | OL generalist | 12099 | 97 | 256 | 0 | 2dpf |
| CP molluscivore | CP generalist | 13768 | 371 | 31 | 0 | 8dpf |
| OL molluscivore | OL generalist | 13694 | 1945 | 443 | 25 | 8dpf |

**Table S4.** Genes differentially expressed between species and misregulated in hybrids that were common to both 8dpf Crescent Pond (CP) and Osprey Lake (OL) comparisons.

| cross | transcript | gene | log2 fold<br>change<br>CP<br>mother vs<br>CP father | log2 fold<br>change OL<br>mother vs<br>OL father | log2 fold<br>change CP<br>parents vs. CP<br>hybrids | log2 fold<br>change OL<br>parents vs. OL<br>hybrids |
| --- | --- | --- | --- | --- | --- | --- |
| generalist × scale-eater | XM_015396529.1 | *trim47* | -1.332 | 0.547 | -1.332 | -1.278 |
| generalist × scale-eater | XM_015405031.1 | *krt13* | -1.184 | -1.181 | -1.183 | -1.229 |
| generalist × scale-eater | XM_015380548.1 | *s100a1* | -1.176 | 0.466 | -1.176 | -0.905 |
| scale-eater ×<br>molluscivore | XM_015396195.1 | *elovl7* | 0.784 | -0.641 | -0.978 | -0.996 |

**Table S5.**
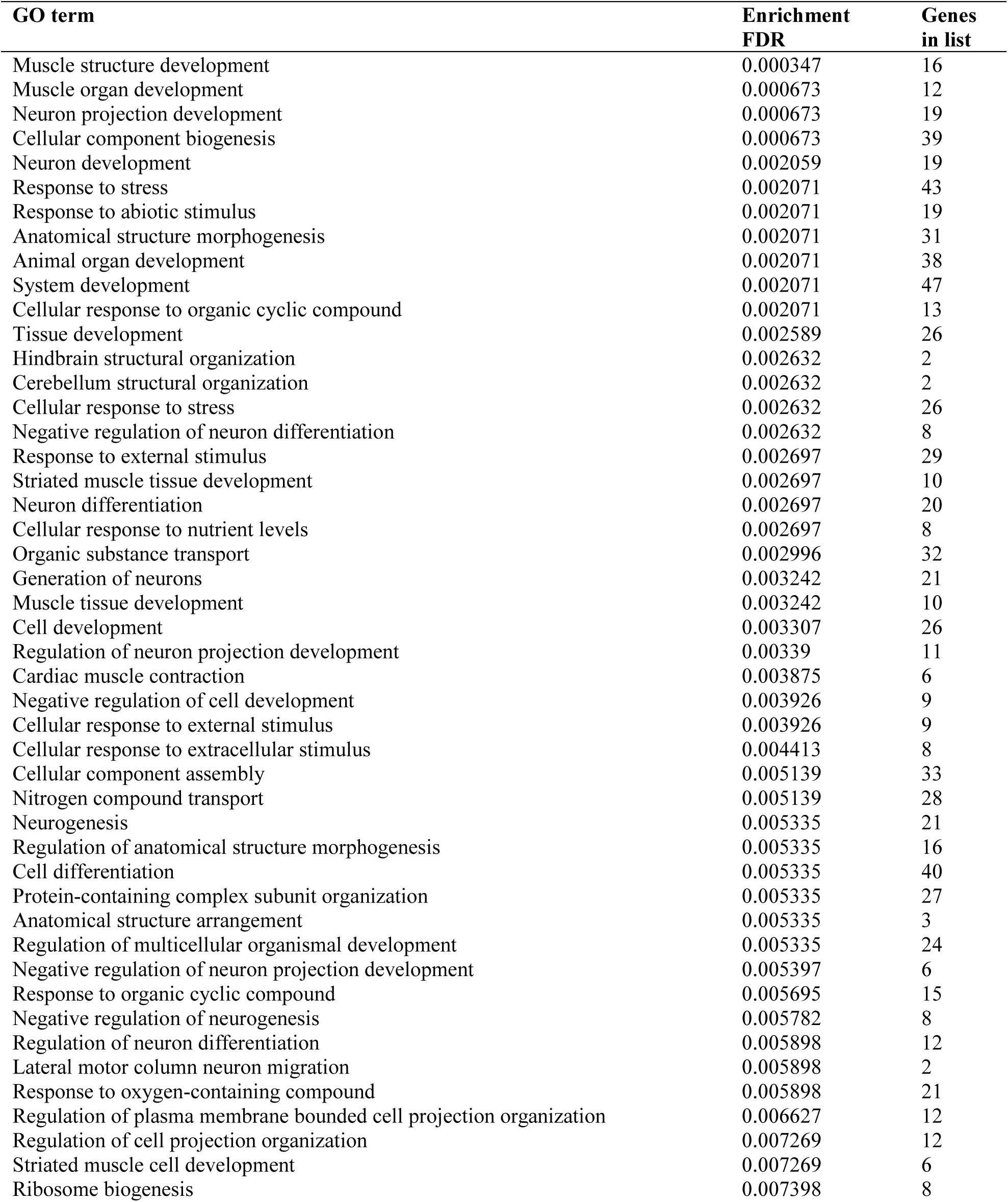
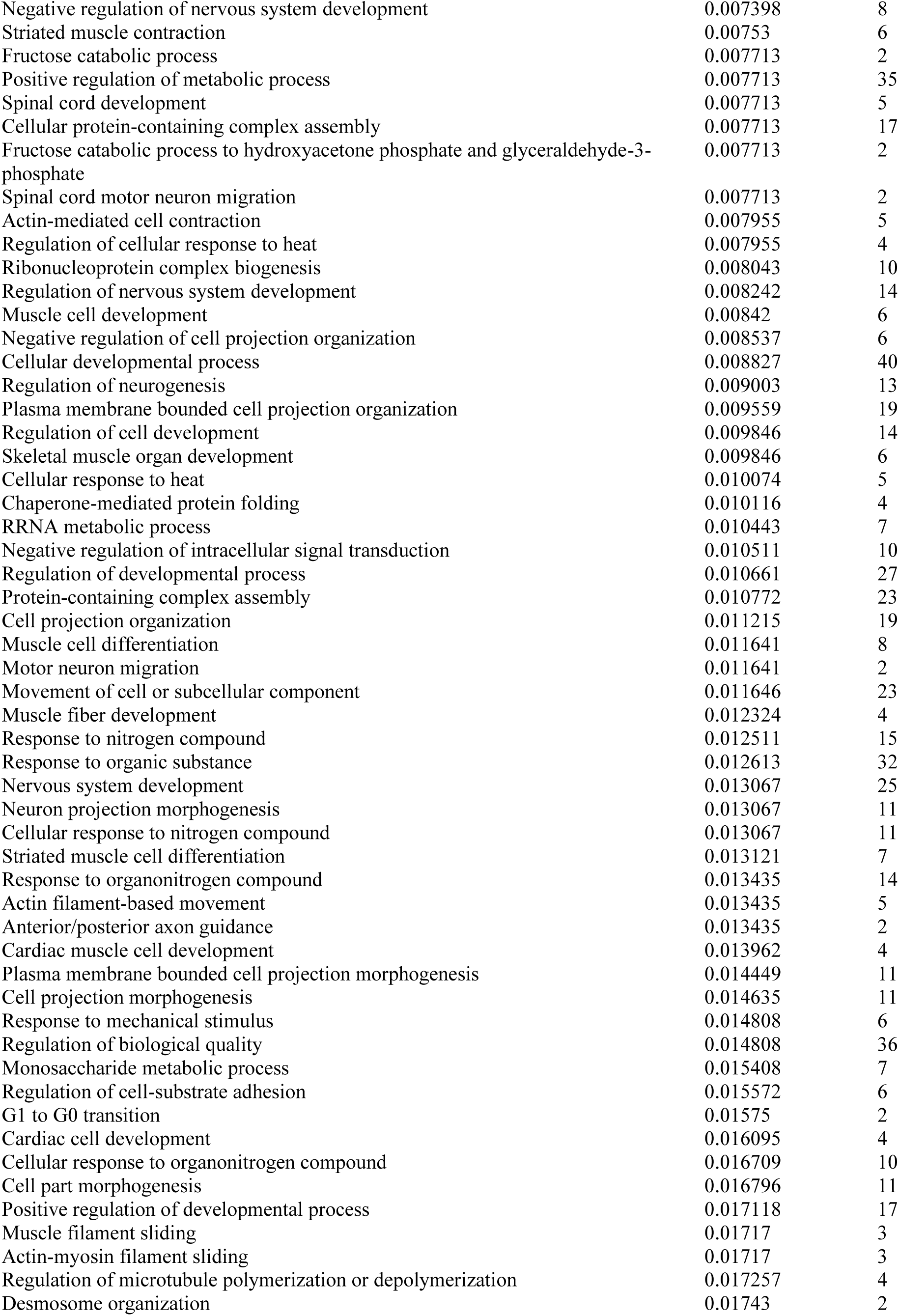
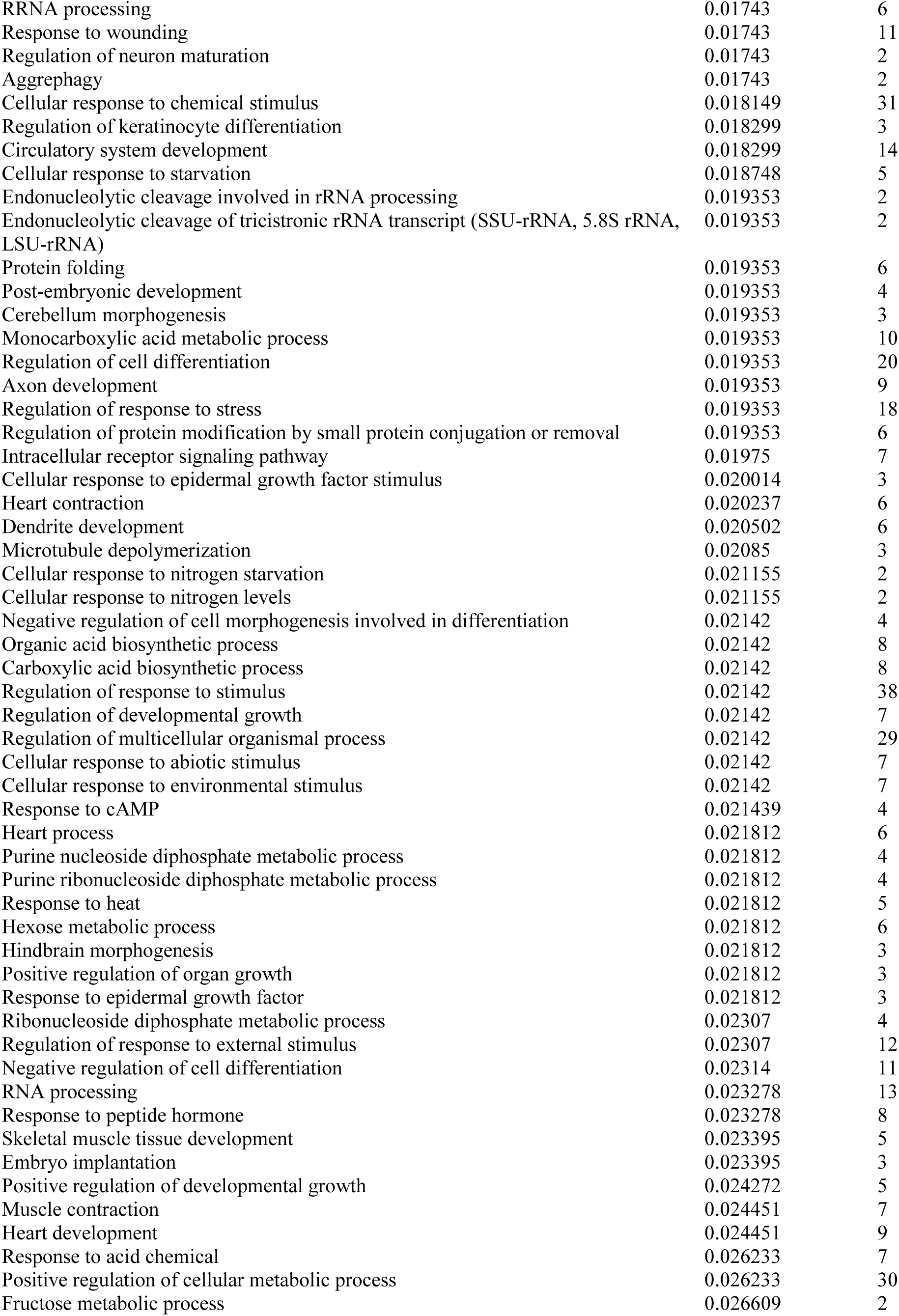
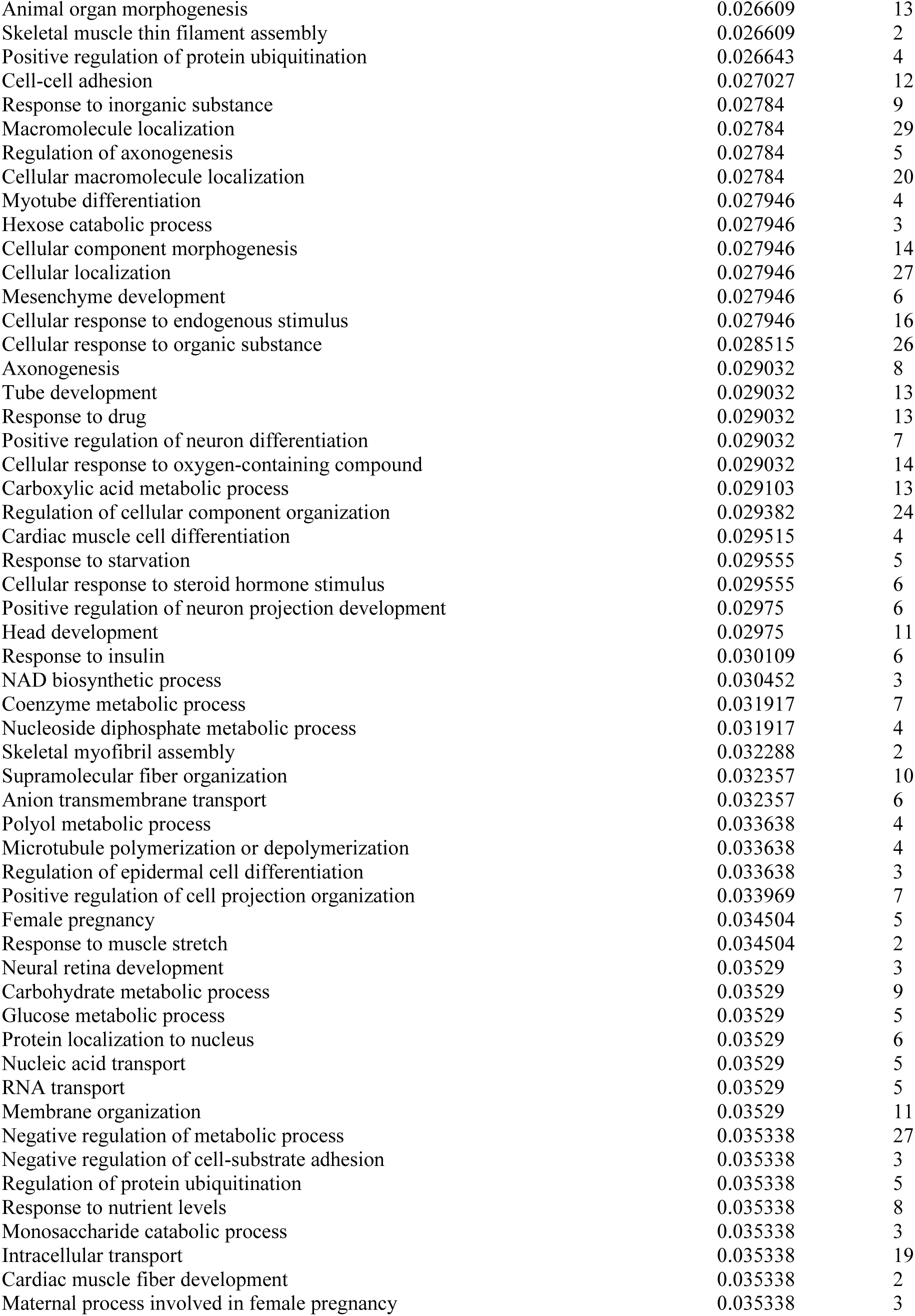
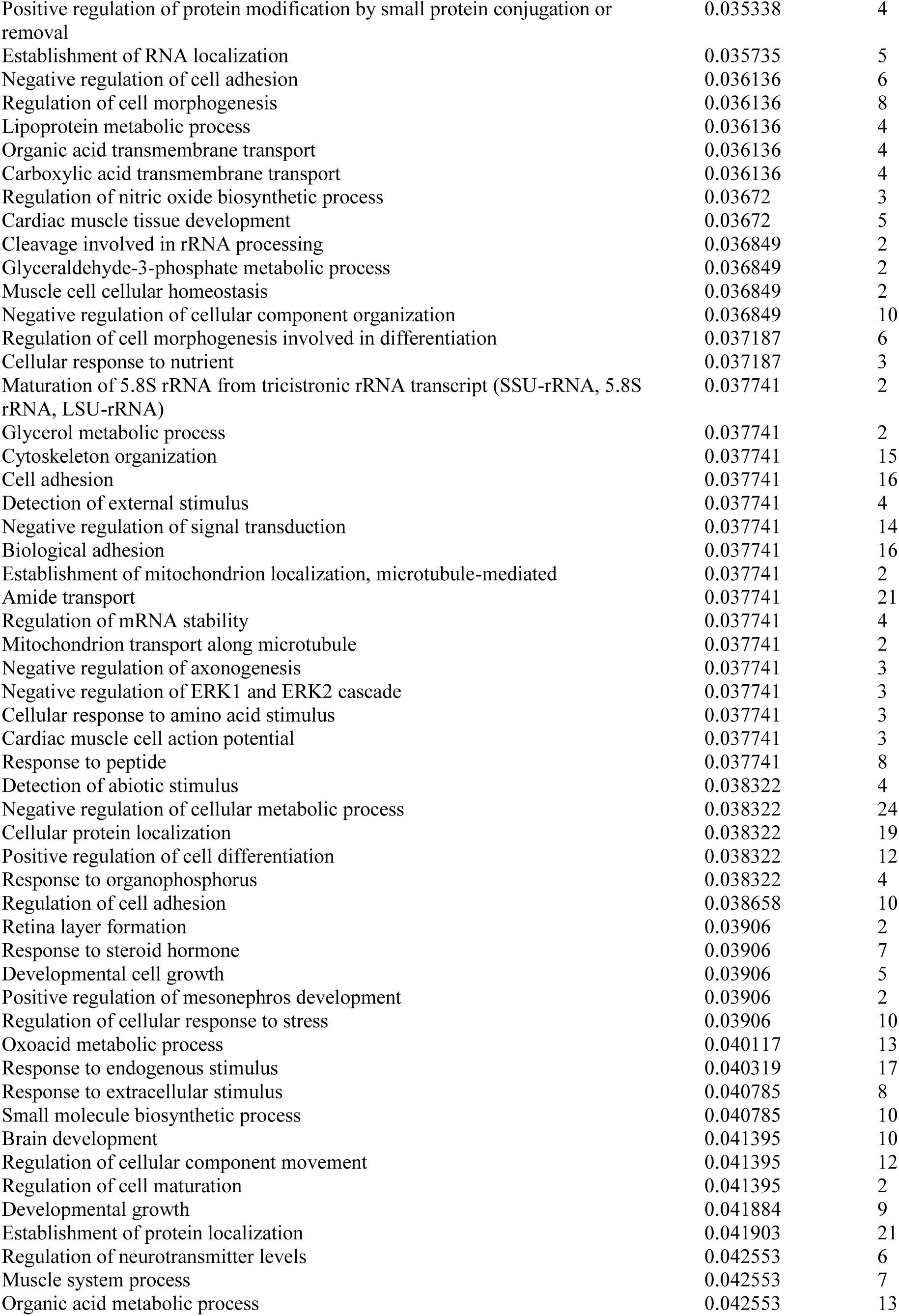
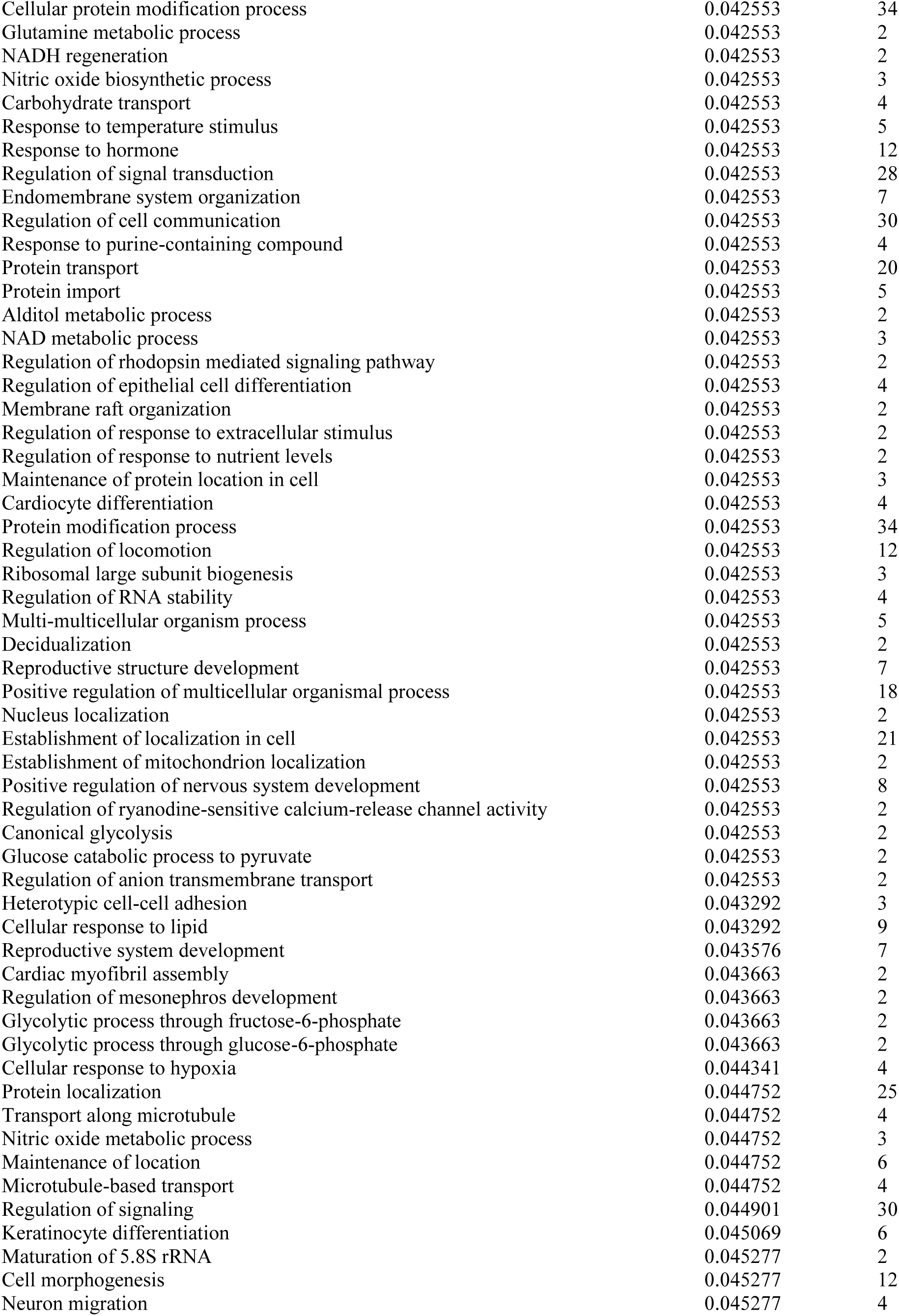
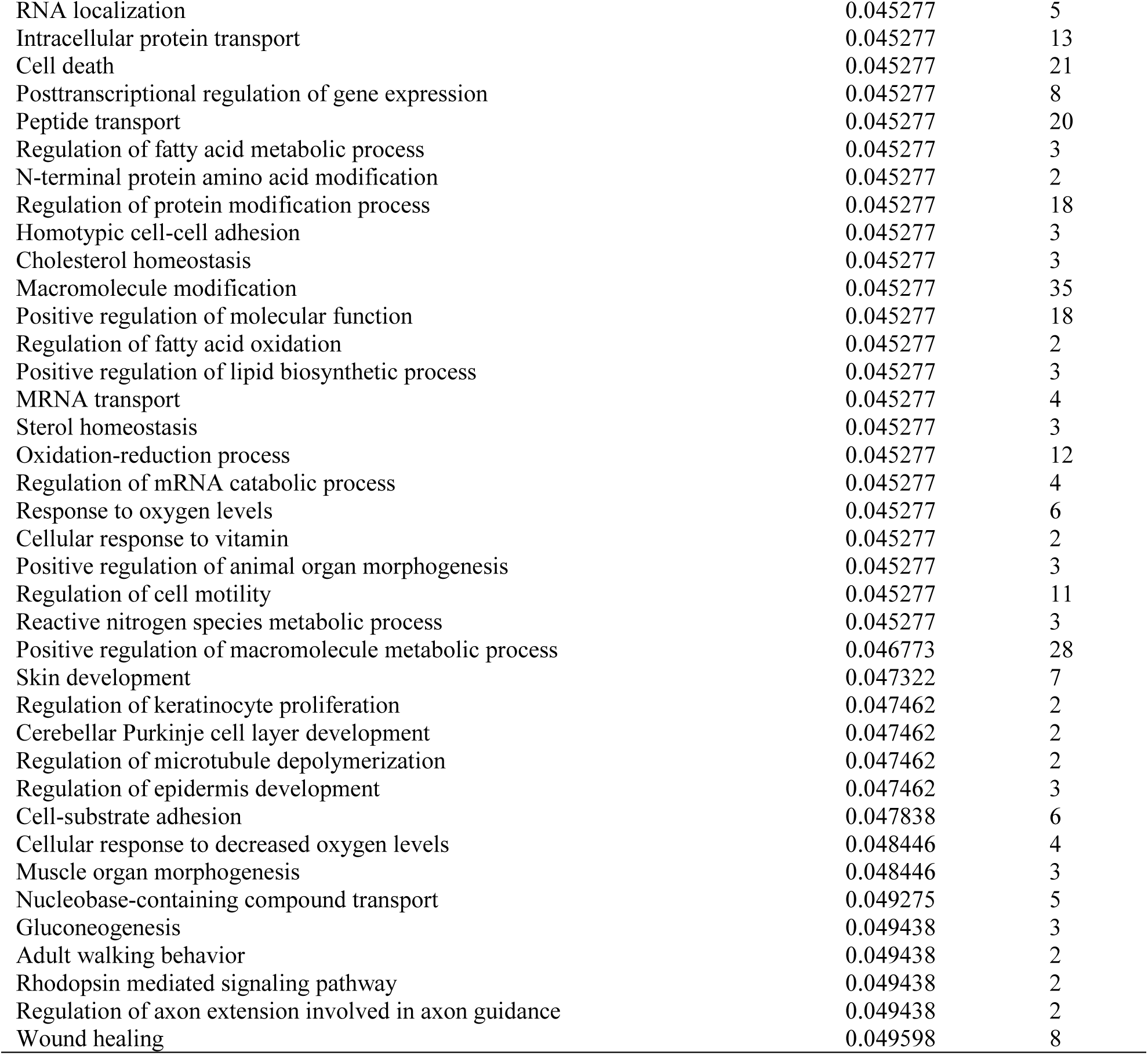
360 significantly enriched gene ontology terms for 125 genes showing differential expression between species and misregulation in F1 hybrids found within highly differentiated regions of the genome.

| GO term | Enrichment<br>FDR | Genes<br>in list |
| --- | --- | --- |
| Muscle structure development | 0.000347 | 16 |
| Muscle organ development | 0.000673 | 12 |
| Neuron projection development | 0.000673 | 19 |
| Cellular component biogenesis | 0.000673 | 39 |
| Neuron development | 0.002059 | 19 |
| Response to stress | 0.002071 | 43 |
| Response to abiotic stimulus | 0.002071 | 19 |
| Anatomical structure morphogenesis | 0.002071 | 31 |
| Animal organ development | 0.002071 | 38 |
| System development | 0.002071 | 47 |
| Cellular response to organic cyclic compound | 0.002071 | 13 |
| Tissue development | 0.002589 | 26 |
| Hindbrain structural organization | 0.002632 | 2 |
| Cerebellum structural organization | 0.002632 | 2 |
| Cellular response to stress | 0.002632 | 26 |
| Negative regulation of neuron differentiation | 0.002632 | 8 |
| Response to external stimulus | 0.002697 | 29 |
| Striated muscle tissue development | 0.002697 | 10 |
| Neuron differentiation | 0.002697 | 20 |
| Cellular response to nutrient levels | 0.002697 | 8 |
| Organic substance transport | 0.002996 | 32 |
| Generation of neurons | 0.003242 | 21 |
| Muscle tissue development | 0.003242 | 10 |
| Cell development | 0.003307 | 26 |
| Regulation of neuron projection development | 0.00339 | 11 |
| Cardiac muscle contraction | 0.003875 | 6 |
| Negative regulation of cell development | 0.003926 | 9 |
| Cellular response to external stimulus | 0.003926 | 9 |
| Cellular response to extracellular stimulus | 0.004413 | 8 |
| Cellular component assembly | 0.005139 | 33 |
| Nitrogen compound transport | 0.005139 | 28 |
| Neurogenesis | 0.005335 | 21 |
| Regulation of anatomical structure morphogenesis | 0.005335 | 16 |
| Cell differentiation | 0.005335 | 40 |
| Protein-containing complex subunit organization | 0.005335 | 27 |
| Anatomical structure arrangement | 0.005335 | 3 |
| Regulation of multicellular organismal development | 0.005335 | 24 |
| Negative regulation of neuron projection development | 0.005397 | 6 |
| Response to organic cyclic compound | 0.005695 | 15 |
| Negative regulation of neurogenesis | 0.005782 | 8 |
| Regulation of neuron differentiation | 0.005898 | 12 |
| Lateral motor column neuron migration | 0.005898 | 2 |
| Response to oxygen-containing compound | 0.005898 | 21 |
| Regulation of plasma membrane bounded cell projection organization | 0.006627 | 12 |
| Regulation of cell projection organization | 0.007269 | 12 |
| Striated muscle cell development | 0.007269 | 6 |
| Ribosome biogenesis | 0.007398 | 8 |
| Negative regulation of nervous system development | 0.007398 | 8 |
| Striated muscle contraction | 0.00753 | 6 |
| Fructose catabolic process | 0.007713 | 2 |
| Positive regulation of metabolic process | 0.007713 | 35 |
| Spinal cord development | 0.007713 | 5 |
| Cellular protein-containing complex assembly | 0.007713 | 17 |
| Fructose catabolic process to hydroxyacetone phosphate and glyceraldehyde-3-phosphate | 0.007713 | 2 |
| Spinal cord motor neuron migration | 0.007713 | 2 |
| Actin-mediated cell contraction | 0.007955 | 5 |
| Regulation of cellular response to heat | 0.007955 | 4 |
| Ribonucleoprotein complex biogenesis | 0.008043 | 10 |
| Regulation of nervous system development | 0.008242 | 14 |
| Muscle cell development | 0.00842 | 6 |
| Negative regulation of cell projection organization | 0.008537 | 6 |
| Cellular developmental process | 0.008827 | 40 |
| Regulation of neurogenesis | 0.009003 | 13 |
| Plasma membrane bounded cell projection organization | 0.009559 | 19 |
| Regulation of cell development | 0.009846 | 14 |
| Skeletal muscle organ development | 0.009846 | 6 |
| Cellular response to heat | 0.010074 | 5 |
| Chaperone-mediated protein folding | 0.010116 | 4 |
| RRNA metabolic process | 0.010443 | 7 |
| Negative regulation of intracellular signal transduction | 0.010511 | 10 |
| Regulation of developmental process | 0.010661 | 27 |
| Protein-containing complex assembly | 0.010772 | 23 |
| Cell projection organization | 0.011215 | 19 |
| Muscle cell differentiation | 0.011641 | 8 |
| Motor neuron migration | 0.011641 | 2 |
| Movement of cell or subcellular component | 0.011646 | 23 |
| Muscle fiber development | 0.012324 | 4 |
| Response to nitrogen compound | 0.012511 | 15 |
| Response to organic substance | 0.012613 | 32 |
| Nervous system development | 0.013067 | 25 |
| Neuron projection morphogenesis | 0.013067 | 11 |
| Cellular response to nitrogen compound | 0.013067 | 11 |
| Striated muscle cell differentiation | 0.013121 | 7 |
| Response to organonitrogen compound | 0.013435 | 14 |
| Actin filament-based movement | 0.013435 | 5 |
| Anterior/posterior axon guidance | 0.013435 | 2 |
| Cardiac muscle cell development | 0.013962 | 4 |
| Plasma membrane bounded cell projection morphogenesis | 0.014449 | 11 |
| Cell projection morphogenesis | 0.014635 | 11 |
| Response to mechanical stimulus | 0.014808 | 6 |
| Regulation of biological quality | 0.014808 | 36 |
| Monosaccharide metabolic process | 0.015408 | 7 |
| Regulation of cell-substrate adhesion | 0.015572 | 6 |
| G1 to G0 transition | 0.01575 | 2 |
| Cardiac cell development | 0.016095 | 4 |
| Cellular response to organonitrogen compound | 0.016709 | 10 |
| Cell part morphogenesis | 0.016796 | 11 |
| Positive regulation of developmental process | 0.017118 | 17 |
| Muscle filament sliding | 0.01717 | 3 |
| Actin-myosin filament sliding | 0.01717 | 3 |
| Regulation of microtubule polymerization or depolymerization | 0.017257 | 4 |
| Desmosome organization | 0.01743 | 2 |
| RRNA processing | 0.01743 | 6 |
| Response to wounding | 0.01743 | 11 |
| Regulation of neuron maturation | 0.01743 | 2 |
| Aggrephagy | 0.01743 | 2 |
| Cellular response to chemical stimulus | 0.018149 | 31 |
| Regulation of keratinocyte differentiation | 0.018299 | 3 |
| Circulatory system development | 0.018299 | 14 |
| Cellular response to starvation | 0.018748 | 5 |
| Endonucleolytic cleavage involved in rRNA processing | 0.019353 | 2 |
| Endonucleolytic cleavage of tricistronic rRNA transcript (SSU-rRNA, 5.8S rRNA, LSU-rRNA) | 0.019353 | 2 |
| Protein folding | 0.019353 | 6 |
| Post-embryonic development | 0.019353 | 4 |
| Cerebellum morphogenesis | 0.019353 | 3 |
| Monocarboxylic acid metabolic process | 0.019353 | 10 |
| Regulation of cell differentiation | 0.019353 | 20 |
| Axon development | 0.019353 | 9 |
| Regulation of response to stress | 0.019353 | 18 |
| Regulation of protein modification by small protein conjugation or removal | 0.019353 | 6 |
| Intracellular receptor signaling pathway | 0.01975 | 7 |
| Cellular response to epidermal growth factor stimulus | 0.020014 | 3 |
| Heart contraction | 0.020237 | 6 |
| Dendrite development | 0.020502 | 6 |
| Microtubule depolymerization | 0.02085 | 3 |
| Cellular response to nitrogen starvation | 0.021155 | 2 |
| Cellular response to nitrogen levels | 0.021155 | 2 |
| Negative regulation of cell morphogenesis involved in differentiation | 0.02142 | 4 |
| Organic acid biosynthetic process | 0.02142 | 8 |
| Carboxylic acid biosynthetic process | 0.02142 | 8 |
| Regulation of response to stimulus | 0.02142 | 38 |
| Regulation of developmental growth | 0.02142 | 7 |
| Regulation of multicellular organismal process | 0.02142 | 29 |
| Cellular response to abiotic stimulus | 0.02142 | 7 |
| Cellular response to environmental stimulus | 0.02142 | 7 |
| Response to cAMP | 0.021439 | 4 |
| Heart process | 0.021812 | 6 |
| Purine nucleoside diphosphate metabolic process | 0.021812 | 4 |
| Purine ribonucleoside diphosphate metabolic process | 0.021812 | 4 |
| Response to heat | 0.021812 | 5 |
| Hexose metabolic process | 0.021812 | 6 |
| Hindbrain morphogenesis | 0.021812 | 3 |
| Positive regulation of organ growth | 0.021812 | 3 |
| Response to epidermal growth factor | 0.021812 | 3 |
| Ribonucleoside diphosphate metabolic process | 0.02307 | 4 |
| Regulation of response to external stimulus | 0.02307 | 12 |
| Negative regulation of cell differentiation | 0.02314 | 11 |
| RNA processing | 0.023278 | 13 |
| Response to peptide hormone | 0.023278 | 8 |
| Skeletal muscle tissue development | 0.023395 | 5 |
| Embryo implantation | 0.023395 | 3 |
| Positive regulation of developmental growth | 0.024272 | 5 |
| Muscle contraction | 0.024451 | 7 |
| Heart development | 0.024451 | 9 |
| Response to acid chemical | 0.026233 | 7 |
| Positive regulation of cellular metabolic process | 0.026233 | 30 |
| Fructose metabolic process | 0.026609 | 2 |
| Animal organ morphogenesis | 0.026609 | 13 |
| Skeletal muscle thin filament assembly | 0.026609 | 2 |
| Positive regulation of protein ubiquitination | 0.026643 | 4 |
| Cell-cell adhesion | 0.027027 | 12 |
| Response to inorganic substance | 0.02784 | 9 |
| Macromolecule localization | 0.02784 | 29 |
| Regulation of axonogenesis | 0.02784 | 5 |
| Cellular macromolecule localization | 0.02784 | 20 |
| Myotube differentiation | 0.027946 | 4 |
| Hexose catabolic process | 0.027946 | 3 |
| Cellular component morphogenesis | 0.027946 | 14 |
| Cellular localization | 0.027946 | 27 |
| Mesenchyme development | 0.027946 | 6 |
| Cellular response to endogenous stimulus | 0.027946 | 16 |
| Cellular response to organic substance | 0.028515 | 26 |
| Axonogenesis | 0.029032 | 8 |
| Tube development | 0.029032 | 13 |
| Response to drug | 0.029032 | 13 |
| Positive regulation of neuron differentiation | 0.029032 | 7 |
| Cellular response to oxygen-containing compound | 0.029032 | 14 |
| Carboxylic acid metabolic process | 0.029103 | 13 |
| Regulation of cellular component organization | 0.029382 | 24 |
| Cardiac muscle cell differentiation | 0.029515 | 4 |
| Response to starvation | 0.029555 | 5 |
| Cellular response to steroid hormone stimulus | 0.029555 | 6 |
| Positive regulation of neuron projection development | 0.02975 | 6 |
| Head development | 0.02975 | 11 |
| Response to insulin | 0.030109 | 6 |
| NAD biosynthetic process | 0.030452 | 3 |
| Coenzyme metabolic process | 0.031917 | 7 |
| Nucleoside diphosphate metabolic process | 0.031917 | 4 |
| Skeletal myofibril assembly | 0.032288 | 2 |
| Supramolecular fiber organization | 0.032357 | 10 |
| Anion transmembrane transport | 0.032357 | 6 |
| Polyol metabolic process | 0.033638 | 4 |
| Microtubule polymerization or depolymerization | 0.033638 | 4 |
| Regulation of epidermal cell differentiation | 0.033638 | 3 |
| Positive regulation of cell projection organization | 0.033969 | 7 |
| Female pregnancy | 0.034504 | 5 |
| Response to muscle stretch | 0.034504 | 2 |
| Neural retina development | 0.03529 | 3 |
| Carbohydrate metabolic process | 0.03529 | 9 |
| Glucose metabolic process | 0.03529 | 5 |
| Protein localization to nucleus | 0.03529 | 6 |
| Nucleic acid transport | 0.03529 | 5 |
| RNA transport | 0.03529 | 5 |
| Membrane organization | 0.03529 | 11 |
| Negative regulation of metabolic process | 0.035338 | 27 |
| Negative regulation of cell-substrate adhesion | 0.035338 | 3 |
| Regulation of protein ubiquitination | 0.035338 | 5 |
| Response to nutrient levels | 0.035338 | 8 |
| Monosaccharide catabolic process | 0.035338 | 3 |
| Intracellular transport | 0.035338 | 19 |
| Cardiac muscle fiber development | 0.035338 | 2 |
| Maternal process involved in female pregnancy | 0.035338 | 3 |
| Positive regulation of protein modification by small protein conjugation or removal | 0.035338 | 4 |
| Establishment of RNA localization | 0.035735 | 5 |
| Negative regulation of cell adhesion | 0.036136 | 6 |
| Regulation of cell morphogenesis | 0.036136 | 8 |
| Lipoprotein metabolic process | 0.036136 | 4 |
| Organic acid transmembrane transport | 0.036136 | 4 |
| Carboxylic acid transmembrane transport | 0.036136 | 4 |
| Regulation of nitric oxide biosynthetic process | 0.03672 | 3 |
| Cardiac muscle tissue development | 0.03672 | 5 |
| Cleavage involved in rRNA processing | 0.036849 | 2 |
| Glyceraldehyde-3-phosphate metabolic process | 0.036849 | 2 |
| Muscle cell cellular homeostasis | 0.036849 | 2 |
| Negative regulation of cellular component organization | 0.036849 | 10 |
| Regulation of cell morphogenesis involved in differentiation | 0.037187 | 6 |
| Cellular response to nutrient | 0.037187 | 3 |
| Maturation of 5.8S rRNA from tricistronic rRNA transcript (SSU-rRNA, 5.8S rRNA, LSU-rRNA) | 0.037741 | 2 |
| Glycerol metabolic process | 0.037741 | 2 |
| Cytoskeleton organization | 0.037741 | 15 |
| Cell adhesion | 0.037741 | 16 |
| Detection of external stimulus | 0.037741 | 4 |
| Negative regulation of signal transduction | 0.037741 | 14 |
| Biological adhesion | 0.037741 | 16 |
| Establishment of mitochondrion localization, microtubule-mediated | 0.037741 | 2 |
| Amide transport | 0.037741 | 21 |
| Regulation of mRNA stability | 0.037741 | 4 |
| Mitochondrion transport along microtubule | 0.037741 | 2 |
| Negative regulation of axonogenesis | 0.037741 | 3 |
| Negative regulation of ERK1 and ERK2 cascade | 0.037741 | 3 |
| Cellular response to amino acid stimulus | 0.037741 | 3 |
| Cardiac muscle cell action potential | 0.037741 | 3 |
| Response to peptide | 0.037741 | 8 |
| Detection of abiotic stimulus | 0.038322 | 4 |
| Negative regulation of cellular metabolic process | 0.038322 | 24 |
| Cellular protein localization | 0.038322 | 19 |
| Positive regulation of cell differentiation | 0.038322 | 12 |
| Response to organophosphorus | 0.038322 | 4 |
| Regulation of cell adhesion | 0.038658 | 10 |
| Retina layer formation | 0.03906 | 2 |
| Response to steroid hormone | 0.03906 | 7 |
| Developmental cell growth | 0.03906 | 5 |
| Positive regulation of mesonephros development | 0.03906 | 2 |
| Regulation of cellular response to stress | 0.03906 | 10 |
| Oxoacid metabolic process | 0.040117 | 13 |
| Response to endogenous stimulus | 0.040319 | 17 |
| Response to extracellular stimulus | 0.040785 | 8 |
| Small molecule biosynthetic process | 0.040785 | 10 |
| Brain development | 0.041395 | 10 |
| Regulation of cellular component movement | 0.041395 | 12 |
| Regulation of cell maturation | 0.041395 | 2 |
| Developmental growth | 0.041884 | 9 |
| Establishment of protein localization | 0.041903 | 21 |
| Regulation of neurotransmitter levels | 0.042553 | 6 |
| Muscle system process | 0.042553 | 7 |
| Organic acid metabolic process | 0.042553 | 13 |
| Cellular protein modification process | 0.042553 | 34 |
| Glutamine metabolic process | 0.042553 | 2 |
| NADH regeneration | 0.042553 | 2 |
| Nitric oxide biosynthetic process | 0.042553 | 3 |
| Carbohydrate transport | 0.042553 | 4 |
| Response to temperature stimulus | 0.042553 | 5 |
| Response to hormone | 0.042553 | 12 |
| Regulation of signal transduction | 0.042553 | 28 |
| Endomembrane system organization | 0.042553 | 7 |
| Regulation of cell communication | 0.042553 | 30 |
| Response to purine-containing compound | 0.042553 | 4 |
| Protein transport | 0.042553 | 20 |
| Protein import | 0.042553 | 5 |
| Alditol metabolic process | 0.042553 | 2 |
| NAD metabolic process | 0.042553 | 3 |
| Regulation of rhodopsin mediated signaling pathway | 0.042553 | 2 |
| Regulation of epithelial cell differentiation | 0.042553 | 4 |
| Membrane raft organization | 0.042553 | 2 |
| Regulation of response to extracellular stimulus | 0.042553 | 2 |
| Regulation of response to nutrient levels | 0.042553 | 2 |
| Maintenance of protein location in cell | 0.042553 | 3 |
| Cardiocyte differentiation | 0.042553 | 4 |
| Protein modification process | 0.042553 | 34 |
| Regulation of locomotion | 0.042553 | 12 |
| Ribosomal large subunit biogenesis | 0.042553 | 3 |
| Regulation of RNA stability | 0.042553 | 4 |
| Multi-multicellular organism process | 0.042553 | 5 |
| Decidualization | 0.042553 | 2 |
| Reproductive structure development | 0.042553 | 7 |
| Positive regulation of multicellular organismal process | 0.042553 | 18 |
| Nucleus localization | 0.042553 | 2 |
| Establishment of localization in cell | 0.042553 | 21 |
| Establishment of mitochondrion localization | 0.042553 | 2 |
| Positive regulation of nervous system development | 0.042553 | 8 |
| Regulation of ryanodine-sensitive calcium-release channel activity | 0.042553 | 2 |
| Canonical glycolysis | 0.042553 | 2 |
| Glucose catabolic process to pyruvate | 0.042553 | 2 |
| Regulation of anion transmembrane transport | 0.042553 | 2 |
| Heterotypic cell-cell adhesion | 0.043292 | 3 |
| Cellular response to lipid | 0.043292 | 9 |
| Reproductive system development | 0.043576 | 7 |
| Cardiac myofibril assembly | 0.043663 | 2 |
| Regulation of mesonephros development | 0.043663 | 2 |
| Glycolytic process through fructose-6-phosphate | 0.043663 | 2 |
| Glycolytic process through glucose-6-phosphate | 0.043663 | 2 |
| Cellular response to hypoxia | 0.044341 | 4 |
| Protein localization | 0.044752 | 25 |
| Transport along microtubule | 0.044752 | 4 |
| Nitric oxide metabolic process | 0.044752 | 3 |
| Maintenance of location | 0.044752 | 6 |
| Microtubule-based transport | 0.044752 | 4 |
| Regulation of signaling | 0.044901 | 30 |
| Keratinocyte differentiation | 0.045069 | 6 |
| Maturation of 5.8S rRNA | 0.045277 | 2 |
| Cell morphogenesis | 0.045277 | 12 |
| Neuron migration | 0.045277 | 4 |
| RNA localization | 0.045277 | 5 |
| Intracellular protein transport | 0.045277 | 13 |
| Cell death | 0.045277 | 21 |
| Posttranscriptional regulation of gene expression | 0.045277 | 8 |
| Peptide transport | 0.045277 | 20 |
| Regulation of fatty acid metabolic process | 0.045277 | 3 |
| N-terminal protein amino acid modification | 0.045277 | 2 |
| Regulation of protein modification process | 0.045277 | 18 |
| Homotypic cell-cell adhesion | 0.045277 | 3 |
| Cholesterol homeostasis | 0.045277 | 3 |
| Macromolecule modification | 0.045277 | 35 |
| Positive regulation of molecular function | 0.045277 | 18 |
| Regulation of fatty acid oxidation | 0.045277 | 2 |
| Positive regulation of lipid biosynthetic process | 0.045277 | 3 |
| MRNA transport | 0.045277 | 4 |
| Sterol homeostasis | 0.045277 | 3 |
| Oxidation-reduction process | 0.045277 | 12 |
| Regulation of mRNA catabolic process | 0.045277 | 4 |
| Response to oxygen levels | 0.045277 | 6 |
| Cellular response to vitamin | 0.045277 | 2 |
| Positive regulation of animal organ morphogenesis | 0.045277 | 3 |
| Regulation of cell motility | 0.045277 | 11 |
| Reactive nitrogen species metabolic process | 0.045277 | 3 |
| Positive regulation of macromolecule metabolic process | 0.046773 | 28 |
| Skin development | 0.047322 | 7 |
| Regulation of keratinocyte proliferation | 0.047462 | 2 |
| Cerebellar Purkinje cell layer development | 0.047462 | 2 |
| Regulation of microtubule depolymerization | 0.047462 | 2 |
| Regulation of epidermis development | 0.047462 | 3 |
| Cell-substrate adhesion | 0.047838 | 6 |
| Cellular response to decreased oxygen levels | 0.048446 | 4 |
| Muscle organ morphogenesis | 0.048446 | 3 |
| Nucleobase-containing compound transport | 0.049275 | 5 |
| Gluconeogenesis | 0.049438 | 3 |
| Adult walking behavior | 0.049438 | 2 |
| Rhodopsin mediated signaling pathway | 0.049438 | 2 |
| Regulation of axon extension involved in axon guidance | 0.049438 | 2 |
| Wound healing | 0.049598 | 8 |

**Table S6.**
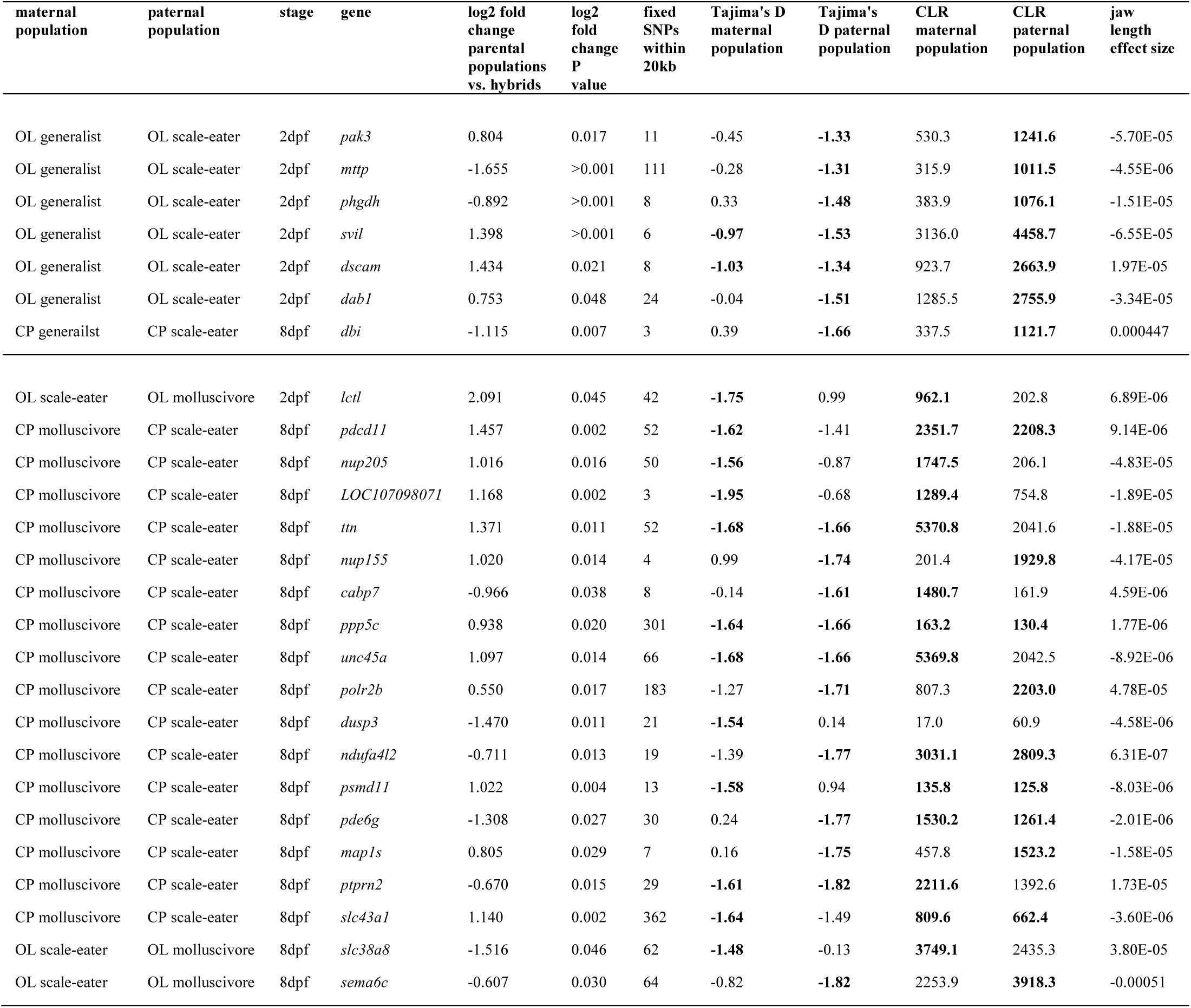
26 genes showing differential expression between species and misregulation in F1 hybrids found within highly differentiated regions of the genome (*F_st_* = 1; *D_xy_* ≥ genome-wide 90^th^ percentile (values in bold; range = 0.0031 – 0.0075; see table S1 for all population thresholds)) that also show strong signs of a hard selective sweep in specialists (negative Tajima’s D < genome-wide 10^th^ percentile (values in bold; range = −1.62 – −0.77 (see Table S7 for all population thresholds); SweeD composite likelihood ratio > 90^th^ percentile for scaffold (values in bold)).

**Table S7.** San Salvador Island population genomic statistics measured across 13.8 million SNPs. Statistics for the top three rows were calculated for all San Salvador individuals of each species (see Fig. S1). The remaining rows are comparisons separated by lake populations used to generate samples for RNAseq (CP = Crescent Pond, OL = Osprey Lake).

| population | mean Tajima's D | Tajima's D 10th percentile | mean $\pi$ |
| --- | --- | --- | --- |
| all generalists | 0.704649 | -0.90273 | 0.003029 |
| all molluscivores | 0.565385 | -1.34112 | 0.002583 |
| all scale-eaters | 0.210182 | -1.62616 | 0.002036 |
| CP generalists | 0.430683 | -1.076 | 0.002806 |
| CP molluscivores | 0.097742 | -1.44811 | 0.00194 |
| CP scale-eaters | -0.01537 | -1.53413 | 0.001385 |
| OL generalists | 0.338391 | -0.77476 | 0.003022 |
| OL molluscivores | 0.227443 | -1.37104 | 0.002458 |
| OL scale-eaters | 0.14957 | -1.31009 | 0.00219 |

**Table S8.**
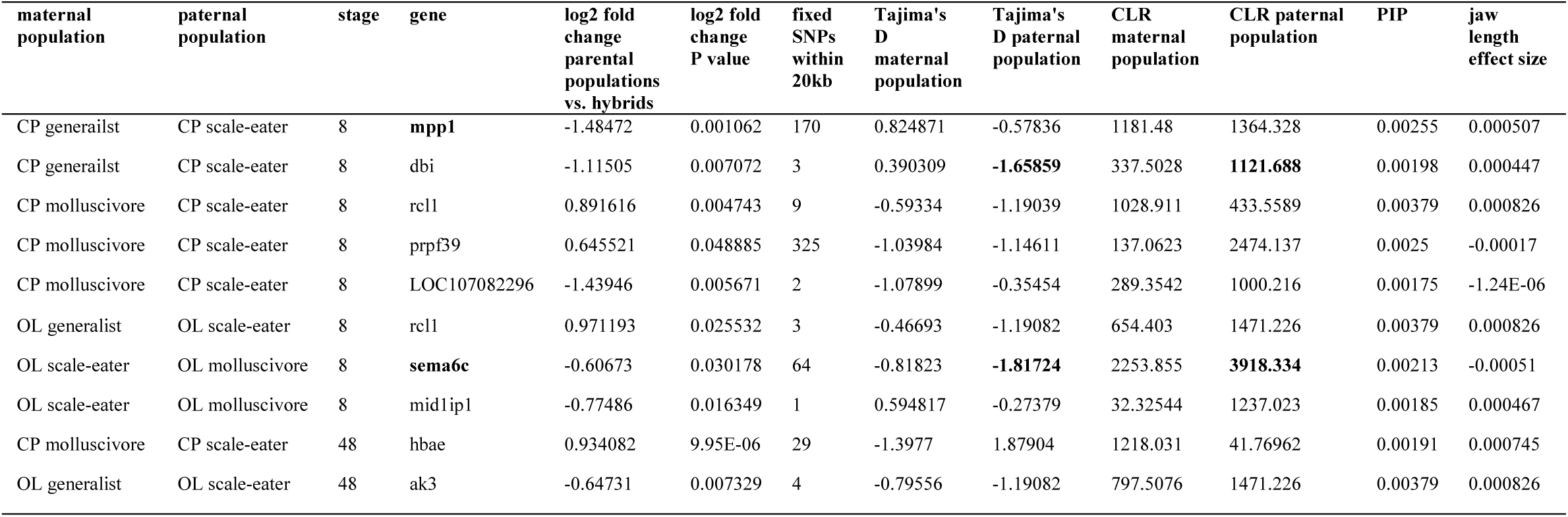
**Ecological DMI candidate genes associated with jaw size.** Nine genes showing differential expression between species and misregulation in F1 hybrids found within highly differentiated regions of the genome (*F_st_* = 1; *D_xy_* ≥ genome-wide 90^th^ percentile (values in bold; range = 0.0075 – 0.0031; see table S1 for all population thresholds)) were also in a 20 kb regions significantly associated with oral jaw size variation across our Caribbean pupfish samples (GEMMA PIP > 99^th^ percentile (0.00175)). Genes in bold are discussed in the main text. The genes *sema6c* and *dbi* (Table S6) also show signs of a hard selective sweep in specialists (negative Tajima’s D < genome-wide 10^th^ percentile; range = −1.62 – −0.77 (see Table S7 for all population thresholds); SweeD composite likelihood ratio > 90^th^ percentile by scaffold (values in bold)).

**Table S9.** Cross design for 124 transcriptomes. All libraries were prepared with Truseq stranded mRNA kits and sequenced at the Vincent J. Coates Genomic Sequencing Center in either May 2018 or June 2018.

| sample ID | stage | sequencing date | parents |
| --- | --- | --- | --- |
| CAE1 | 8dpf | May-18 | Crescent Pond generalists |
| CAE2 | 8dpf | May-18 | Crescent Pond generalists |
| CAE3 | 8dpf | May-18 | Crescent Pond generalists |
| CAE4 | 8dpf | May-18 | Crescent Pond generalists |
| CAE5 | 8dpf | May-18 | Crescent Pond generalists |
| CME1 | 8dpf | Jul-18 | Crescent Pond snail eaters |
| CME2 | 8dpf | Jul-18 | Crescent Pond snail eaters |
| CME5 | 8dpf | Jul-18 | Crescent Pond snail eaters |
| CPE1 | 8dpf | May-18 | Crescent Pond scale eaters |
| CPE2 | 8dpf | May-18 | Crescent Pond scale eaters |
| CPE3 | 8dpf | May-18 | Crescent Pond scale eaters |
| CPE4 | 8dpf | May-18 | Crescent Pond scale eaters |
| CPE5 | 8dpf | May-18 | Crescent Pond scale eaters |
| CQE1 | 8dpf | Jul-18 | New Providence female x New Providence generalist male |
| CQE2 | 8dpf | Jul-18 | New Providence female x New Providence generalist male |
| CQE3 | 8dpf | Jul-18 | New Providence female x New Providence generalist male |
| NCE1 | 8dpf | May-18 | North Carolina generalists |
| NCE2 | 8dpf | May-18 | North Carolina generalists |
| NCE3 | 8dpf | May-18 | North Carolina generalists |
| NCE4 | 8dpf | May-18 | North Carolina generalists |
| NCE5 | 8dpf | May-18 | North Carolina generalists |
| OAE1 | 8dpf | May-18 | Osprey Lake generalists |
| OAE2 | 8dpf | May-18 | Osprey Lake generalists |
| OAE3 | 8dpf | May-18 | Osprey Lake generalists |
| OAE4 | 8dpf | May-18 | Osprey Lake generalists |
| OME1 | 8dpf | May-18 | Osprey Lake snail eaters |
| OME2 | 8dpf | May-18 | Osprey Lake snail eaters |
| OME3 | 8dpf | May-18 | Osprey Lake snail eaters |
| OME4 | 8dpf | May-18 | Osprey Lake snail eaters |
| OME5 | 8dpf | May-18 | Osprey Lake snail eaters |
| OPE1 | 8dpf | May-18 | Osprey Lake scale eaters |
| OPE2 | 8dpf | May-18 | Osprey Lake scale eaters |
| OPE3 | 8dpf | May-18 | Osprey Lake scale eaters |
| OPE4 | 8dpf | May-18 | Osprey Lake scale eaters |
| OPE5 | 8dpf | May-18 | Osprey Lake scale eaters |
| CPU1 | 8dpf | Jul-18 | Crescent Pond generalist female x Crescent Pond snail eater male |
| CPU3 | 8dpf | Jul-18 | Crescent Pond generalist female x Crescent Pond snail eater male |
| CPU5 | 8dpf | Jul-18 | Crescent Pond generalist female x Crescent Pond snail eater male |
| CVE1 | 8dpf | Jul-18 | Crescent Pond generalist female x Crescent Pond scale eater male |
| CVE2 | 8dpf | Jul-18 | Crescent Pond generalist female x Crescent Pond scale eater male |
| CVE5 | 8dpf | Jul-18 | Crescent Pond generalist female x Crescent Pond scale eater male |
| CWE2 | 8dpf | Jul-18 | Crescent Pond snail eater female x Crescent Pond scale eater male |
| CWE3 | 8dpf | Jul-18 | Crescent Pond snail eater female x Crescent Pond scale eater male |
| CWE4 | 8dpf | Jul-18 | Crescent Pond snail eater female x Crescent Pond scale eater male |
| CXE2 | 8dpf | Jul-18 | Crescent Pond snail eater female x Crescent Pond generalist male |
| CXE3 | 8dpf | Jul-18 | Crescent Pond snail eater female x Crescent Pond generalist male |
| CXE4 | 8dpf | Jul-18 | Crescent Pond snail eater female x Crescent Pond generalist male |
| NAE1 | 8dpf | Jul-18 | North Carolina female x Crescent Pond generalist male |
| NAE2 | 8dpf | Jul-18 | North Carolina female x Crescent Pond generalist male |
| NAE4 | 8dpf | Jul-18 | North Carolina female x Crescent Pond generalist male |
| OUE1 | 8dpf | Jul-18 | Osprey Lake generalist female x Osprey Lake snail eater male |
| OUE3 | 8dpf | Jul-18 | Osprey Lake generalist female x Osprey Lake snail eater male |
| OUE4 | 8dpf | Jul-18 | Osprey Lake generalist female x Osprey Lake snail eater male |
| OVE1 | 8dpf | Jul-18 | Osprey Lake generalist female x Osprey Lake scale eater male |
| OVE4 | 8dpf | Jul-18 | Osprey Lake generalist female x Osprey Lake scale eater male |
| OVE5 | 8dpf | Jul-18 | Osprey Lake generalist female x Osprey Lake scale eater male |
| OXE2 | 8dpf | Jul-18 | Osprey Lake snail eater female x Osprey Lake generalist male |
| OYE1 | 8dpf | May-18 | Osprey Lake scale eater female x Osprey Lake generalist male |
| OYE2 | 8dpf | May-18 | Osprey Lake scale eater female x Osprey Lake generalist male |
| OYE3 | 8dpf | May-18 | Osprey Lake scale eater female x Osprey Lake generalist male |
| OYE4 | 8dpf | May-18 | Osprey Lake scale eater female x Osprey Lake generalist male |
| OYE5 | 8dpf | May-18 | Osprey Lake scale eater female x Osprey Lake generalist male |
| OZE2 | 8dpf | Jul-18 | Osprey Lake scale eater female x Osprey Lake snail eater male |
| OZE4 | 8dpf | Jul-18 | Osprey Lake scale eater female x Osprey Lake snail eater male |
| OZE5 | 8dpf | Jul-18 | Osprey Lake scale eater female x Osprey Lake snail eater male |
| PAE1 | 8dpf | Jul-18 | New Providence female x Crescent Pond generalist |
| PAE2 | 8dpf | Jul-18 | New Providence female x Crescent Pond generalist |
| PAE5 | 8dpf | Jul-18 | New Providence female x Crescent Pond generalist |
| CAT1 | 2dpf | May-18 | Crescent Pond generalists |
| CAT2 | 2dpf | May-18 | Crescent Pond generalists |
| CAT3 | 2dpf | May-18 | Crescent Pond generalists |
| CMT1 | 2dpf | Jul-18 | Crescent Pond snail eaters |
| CMT2 | 2dpf | Jul-18 | Crescent Pond snail eaters |
| CMT3 | 2dpf | Jul-18 | Crescent Pond snail eaters |
| CPT1 | 2dpf | May-18 | Crescent Pond scale eaters |
| CPT2 | 2dpf | May-18 | Crescent Pond scale eaters |
| CPT3 | 2dpf | Jul-18 | Crescent Pond scale eaters |
| CQT1 | 2dpf | Jul-18 | New Providence female x New Providence generalist male |
| CQT2 | 2dpf | Jul-18 | New Providence female x New Providence generalist male |
| NCT1 | 2dpf | May-18 | North Carolina generalists |
| NCT2 | 2dpf | May-18 | North Carolina generalists |
| NCT3 | 2dpf | May-18 | North Carolina generalists |
| OAT1 | 2dpf | May-18 | Osprey Lake generalists |
| OAT2 | 2dpf | May-18 | Osprey Lake generalists |
| OAT3 | 2dpf | Jul-18 | Osprey Lake generalists |
| OMT1 | 2dpf | May-18 | Osprey Lake snail eaters |
| OMT2 | 2dpf | May-18 | Osprey Lake snail eaters |
| OMT3 | 2dpf | May-18 | Osprey Lake snail eaters |
| OPT1 | 2dpf | May-18 | Osprey Lake scale eaters |
| OPT2 | 2dpf | May-18 | Osprey Lake scale eaters |
| OPT3 | 2dpf | May-18 | Osprey Lake scale eaters |
| CUT1 | 2dpf | Jul-18 | Crescent Pond generalist female x Crescent Pond snail eater male |
| CUT2 | 2dpf | Jul-18 | Crescent Pond generalist female x Crescent Pond snail eater male |
| CUT3 | 2dpf | Jul-18 | Crescent Pond generalist female x Crescent Pond snail eater male |
| CVT1 | 2dpf | May-18 | Crescent Pond generalist female x Crescent Pond scale eater male |
| CVT2 | 2dpf | May-18 | Crescent Pond generalist female x Crescent Pond scale eater male |
| CVT3 | 2dpf | May-18 | Crescent Pond generalist female x Crescent Pond scale eater male |
| CWT1 | 2dpf | May-18 | Crescent Pond snail eater female x Crescent Pond scale eater male |
| CWT2 | 2dpf | May-18 | Crescent Pond snail eater female x Crescent Pond scale eater male |
| CWT3 | 2dpf | May-18 | Crescent Pond snail eater female x Crescent Pond scale eater male |
| CXT1 | 2dpf | May-18 | Crescent Pond snail eater female x Crescent Pond generalist male |
| CXT2 | 2dpf | May-18 | Crescent Pond snail eater female x Crescent Pond generalist male |
| CXT3 | 2dpf | May-18 | Crescent Pond snail eater female x Crescent Pond generalist male |
| NAT1 | 2dpf | May-18 | North Carolina female x Crescent Pond generalist male |
| NAT2 | 2dpf | May-18 | North Carolina female x Crescent Pond generalist male |
| NAT3 | 2dpf | May-18 | North Carolina female x Crescent Pond generalist male |
| OUT1 | 2dpf | Jul-18 | Osprey Lake generalist female x Osprey Lake snail eater male |
| OUT2 | 2dpf | Jul-18 | Osprey Lake generalist female x Osprey Lake snail eater male |
| OUT3 | 2dpf | Jul-18 | Osprey Lake generalist female x Osprey Lake snail eater male |
| OVT1 | 2dpf | May-18 | Osprey Lake generalist female x Osprey Lake scale eater male |
| OVT2 | 2dpf | May-18 | Osprey Lake generalist female x Osprey Lake scale eater male |
| OVT3 | 2dpf | May-18 | Osprey Lake generalist female x Osprey Lake scale eater male |
| OXT1 | 2dpf | Jul-18 | Osprey Lake snail eater female x Osprey Lake generalist male |
| OXT2 | 2dpf | Jul-18 | Osprey Lake snail eater female x Osprey Lake generalist male |
| OXT3 | 2dpf | Jul-18 | Osprey Lake snail eater female x Osprey Lake generalist male |
| OYT1 | 2dpf | May-18 | Osprey Lake scale eater female x Osprey Lake generalist male |
| OYT2 | 2dpf | May-18 | Osprey Lake scale eater female x Osprey Lake generalist male |
| OYT3 | 2dpf | May-18 | Osprey Lake scale eater female x Osprey Lake generalist male |
| OZT1 | 2dpf | Jul-18 | Osprey Lake scale eater female x Osprey Lake snail eater male |
| OZT2 | 2dpf | Jul-18 | Osprey Lake scale eater female x Osprey Lake snail eater male |
| OZT3 | 2dpf | Jul-18 | Osprey Lake scale eater female x Osprey Lake snail eater male |
| PAT1 | 2dpf | May-18 | New Providence female x Crescent Pond generalist |
| PAT2 | 2dpf | May-18 | New Providence female x Crescent Pond generalist |
| PAT3 | 2dpf | May-18 | New Providence female x Crescent Pond generalist |

**Fig S1.**
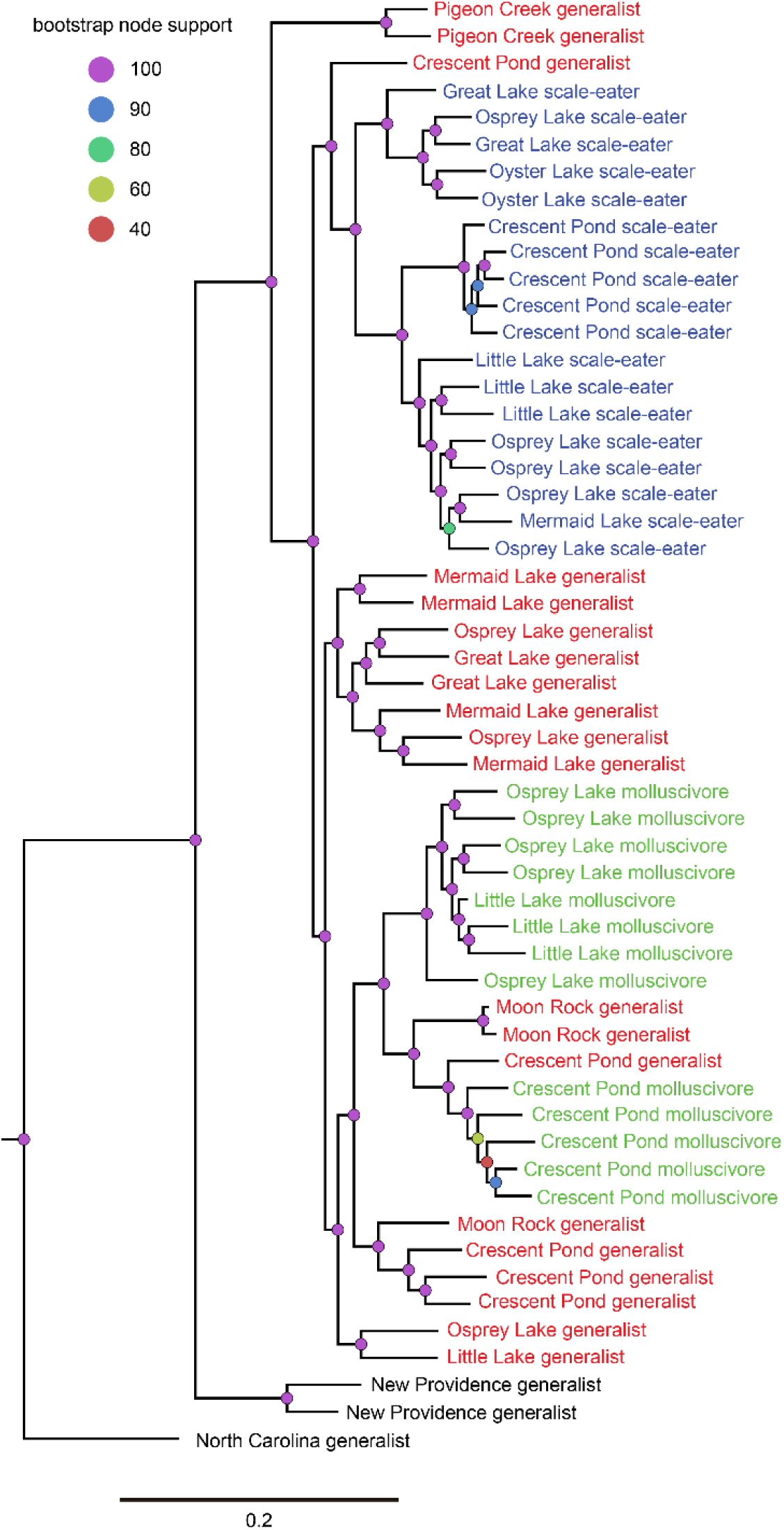
Maximum likelihood tree generated using RAxML with 1.7 million SNPs showing phylogenetic relationships between 55 *Cyprinodon* individuals. Relationships for three outgroup individuals that were included in the genomic dataset are not shown. Red = San Salvador generalist, green = molluscivore, blue = scale-eater, black = outgroup generalist.

**Fig S2.**
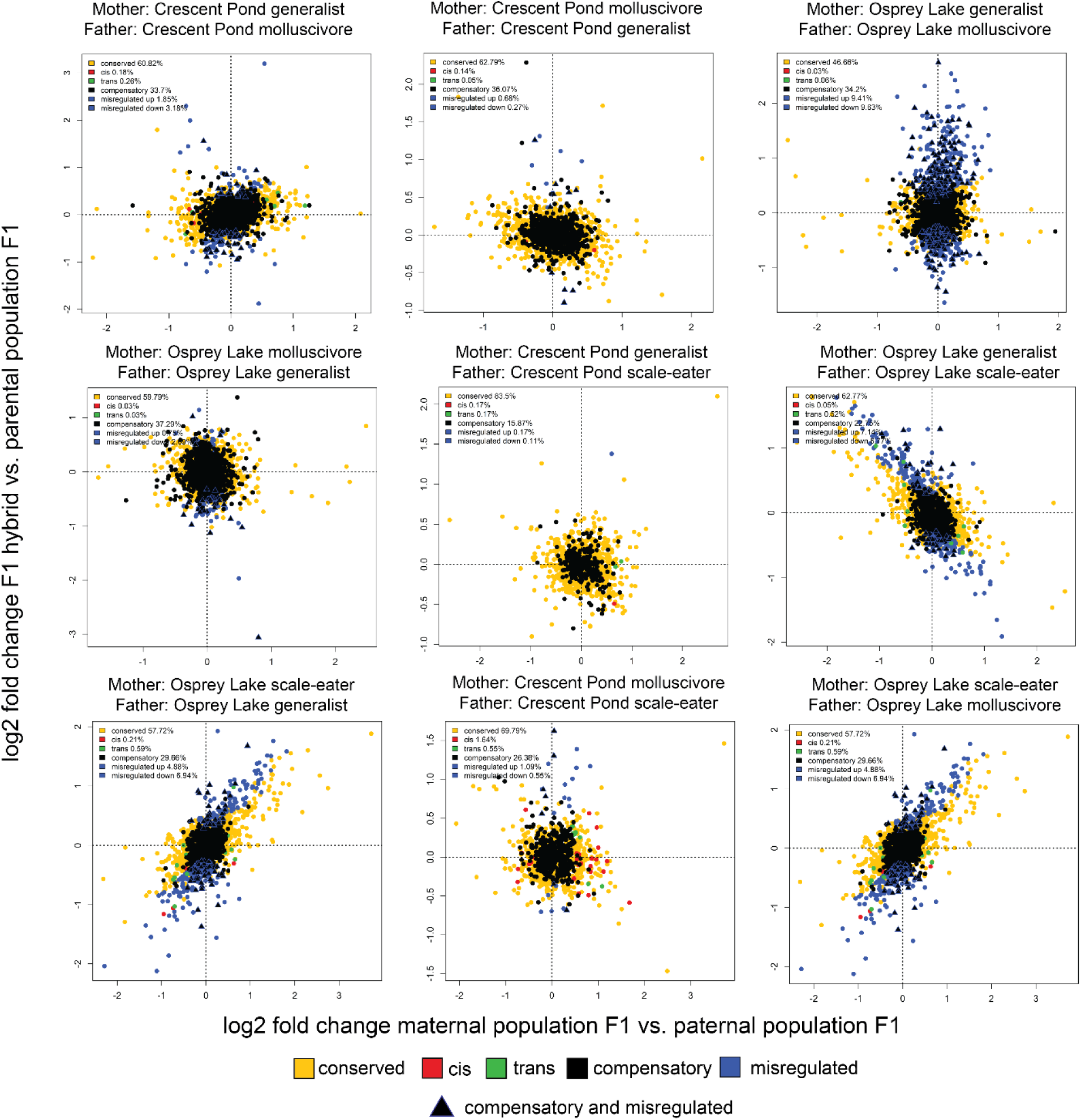
Regulatory mechanisms underlying expression divergence at 2 dpf in San Salvador crosses. Yellow = conserved (no difference in expression between any group or ambiguous expression patterns), red = *cis* (significant ASE in hybrids, significant differential expression between parental populations of purebred F1 offspring, and no significant *trans*-contribution), green = *trans* (significant ASE in hybrids, significant differential expression between parental populations of purebred F1 offspring, and significant *trans*-contribution), black = compensatory (significant ASE in hybrids, no significant differential expression between parental populations of purebred F1 offspring), blue = misregulated (significant differential expression between purebred F1 and hybrid F1), triangle = compensatory and misregulated.

**Fig S3.**
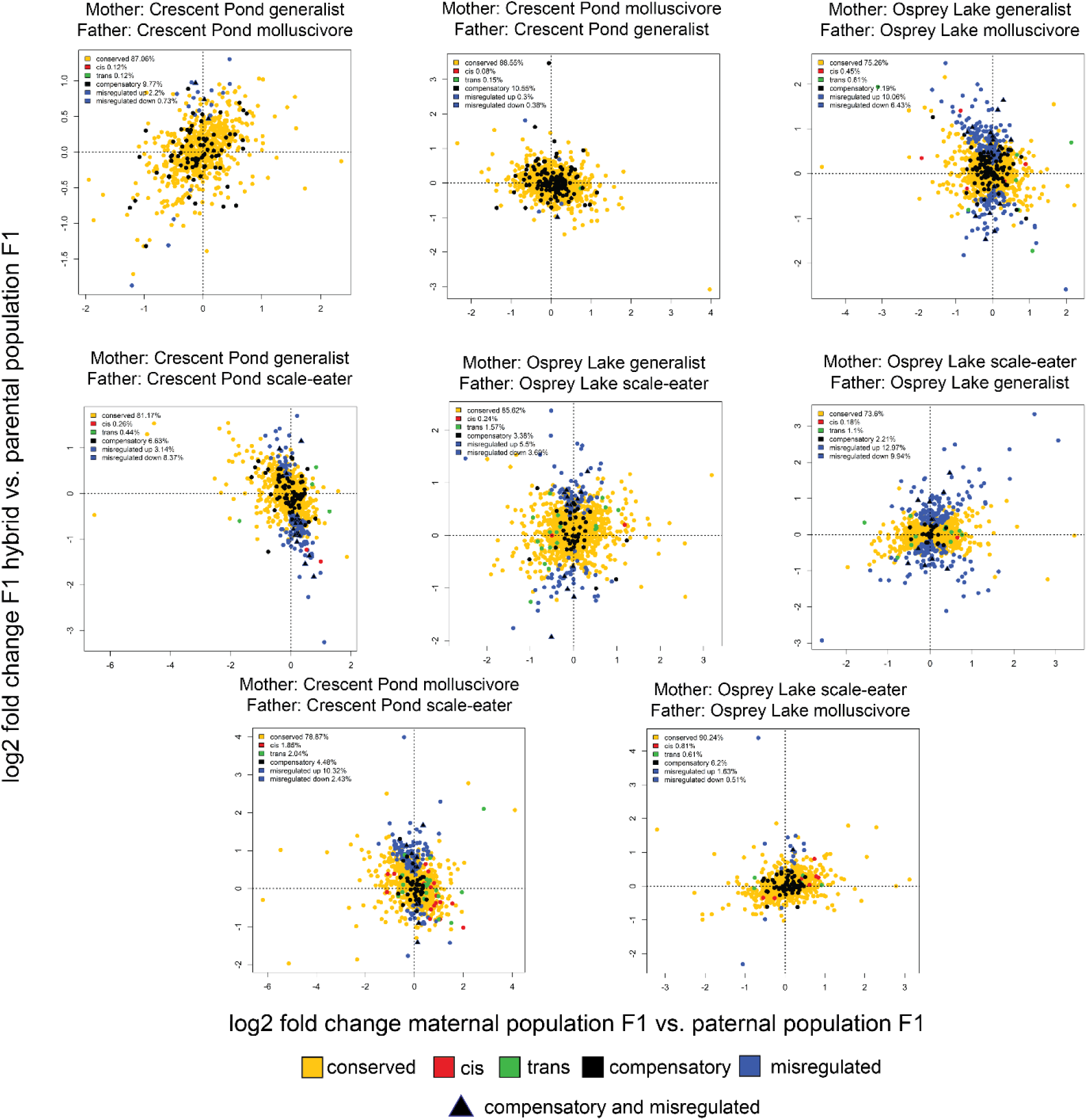
Regulatory mechanisms underlying expression divergence at 8 dpf in San Salvador crosses. Yellow = conserved (no difference in expression between any group or ambiguous expression patterns), red = *cis* (significant ASE in hybrids, significant differential expression between parental populations of purebred F1 offspring, and no significant *trans*-contribution), green = *trans* (significant ASE in hybrids, significant differential expression between parental populations of purebred F1 offspring, and significant *trans*-contribution), black = compensatory (significant ASE in hybrids, no significant differential expression between parental populations of purebred F1 offspring), blue = misregulated (significant differential expression between purebred F1 and hybrid F1), triangle = compensatory and misregulated.

**Fig S4.**
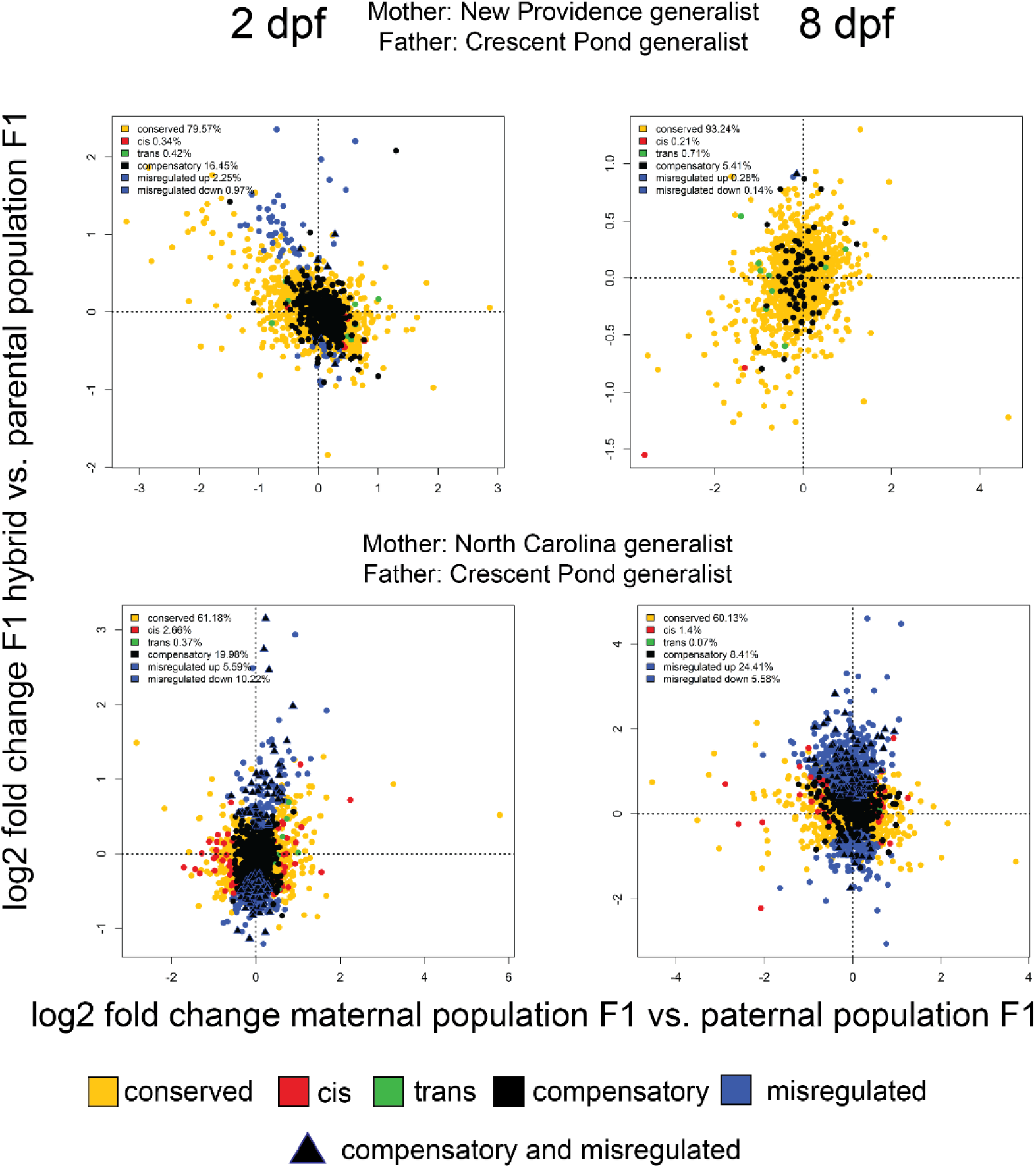
Regulatory mechanisms underlying expression divergence in outgroup generalist population crosses. Yellow = conserved (no difference in expression between any group or ambiguous expression patterns), red = *cis* (significant ASE in hybrids, significant differential expression between parental populations of purebred F1 offspring, and no significant *trans*-contribution), green = *trans* (significant ASE in hybrids, significant differential expression between parental populations of purebred F1 offspring, and significant *trans*-contribution), black = compensatory (significant ASE in hybrids, no significant differential expression between parental populations of purebred F1 offspring), blue = misregulated (significant differential expression between purebred F1 and hybrid F1), triangle = compensatory and misregulated.

**Fig. S5.**
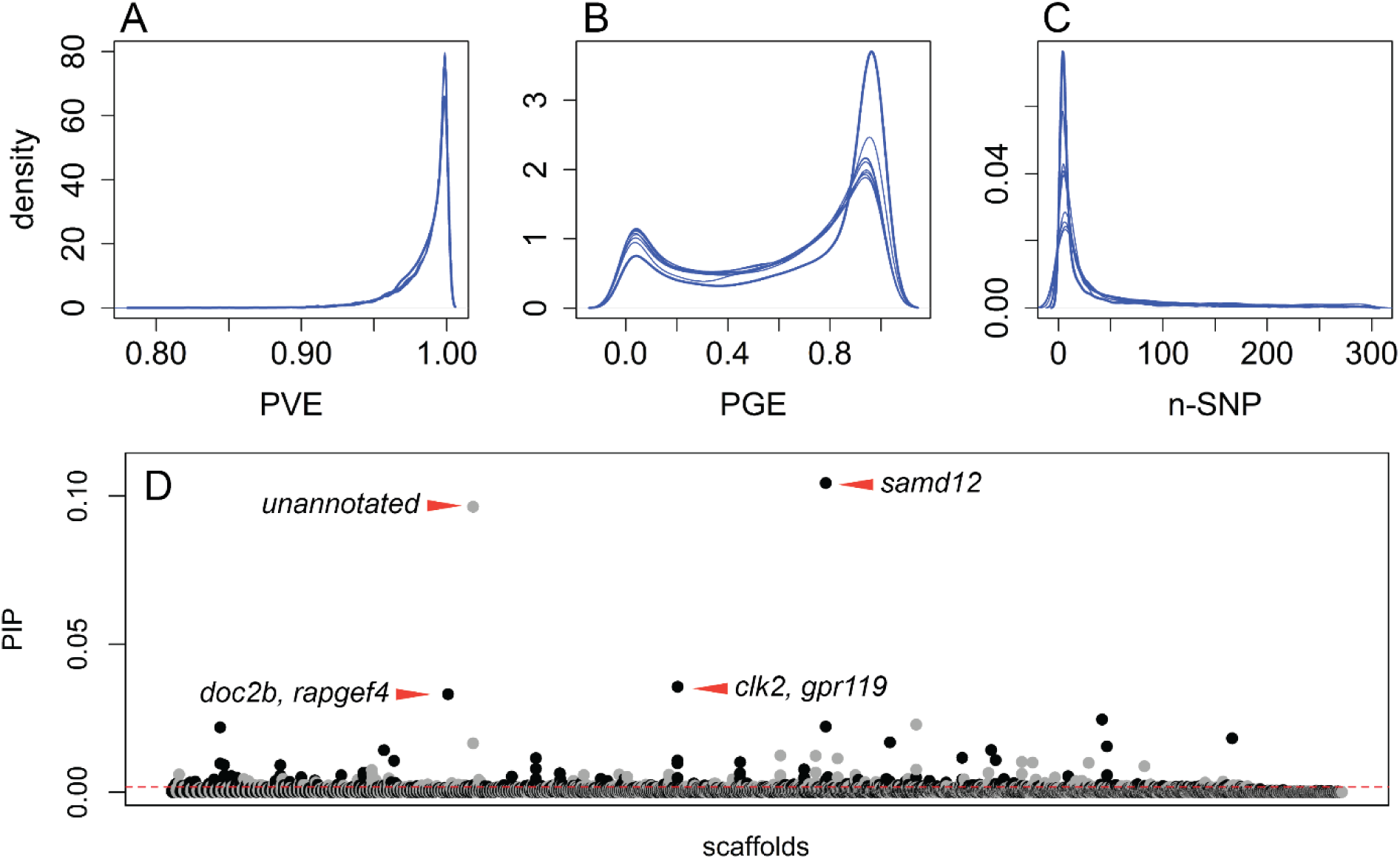
Genome-wide association mapping. GEMMA implements a Bayesian sparse linear mixed model (BSLMM) that uses MCMC to estimate the proportion of phenotypic variation explained by every SNP included in the analysis (A; PVE), the proportion of phenotypic variation explained by SNPs of large effect (B; PGE), which are defined as SNPs with a non-zero effect on the phenotype, and the number of large-effect SNPs needed to explain PGE (C; nSNPs). Each blue line represents one of ten independent runs of the BSLMM. D) Posterior inclusion probability for 20 kb windows across all scaffolds (alternating black and grey for each scaffold). Windows that showed PIP values above the 99th percentile (0.00175; dotted red line) were considered to have a significant effect on jaw size variation. Red arrows indicate genes within top four windows (*samd12, clk2, gpr119, doc2b, rapgef4*).

**Fig S6.**
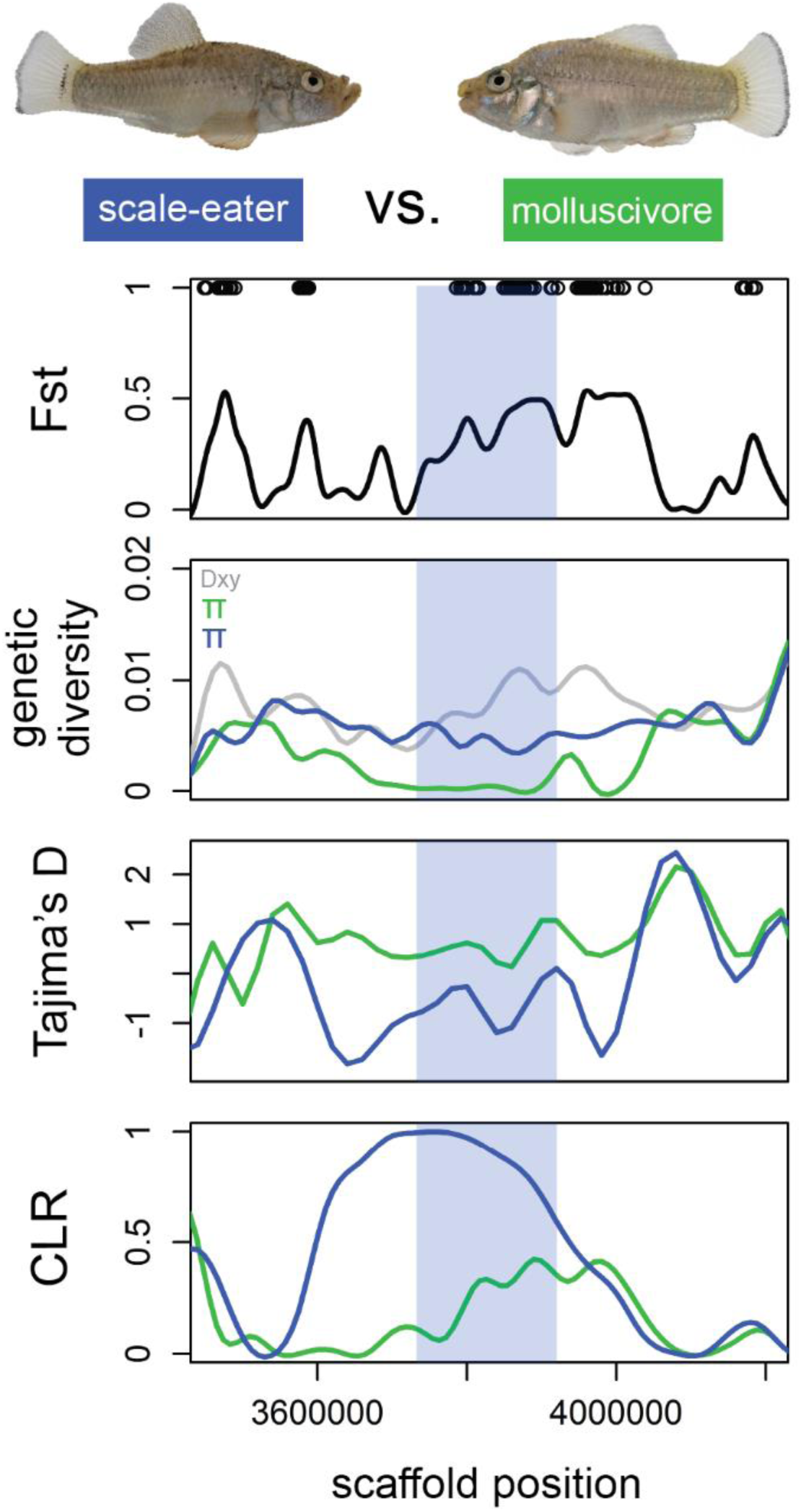
The *sema6c* gene region (light blue) contains 64 SNPs fixed between Osprey Lake scale-eaters (blue) vs. molluscivores (green), shows strong between-population divergence and low within-population diversity, shows strong signs of a hard selective sweep, and is significantly associated with oral jaw length variation in a genome-wide association analysis using GEMMA (Table S8).

**Fig S7.**
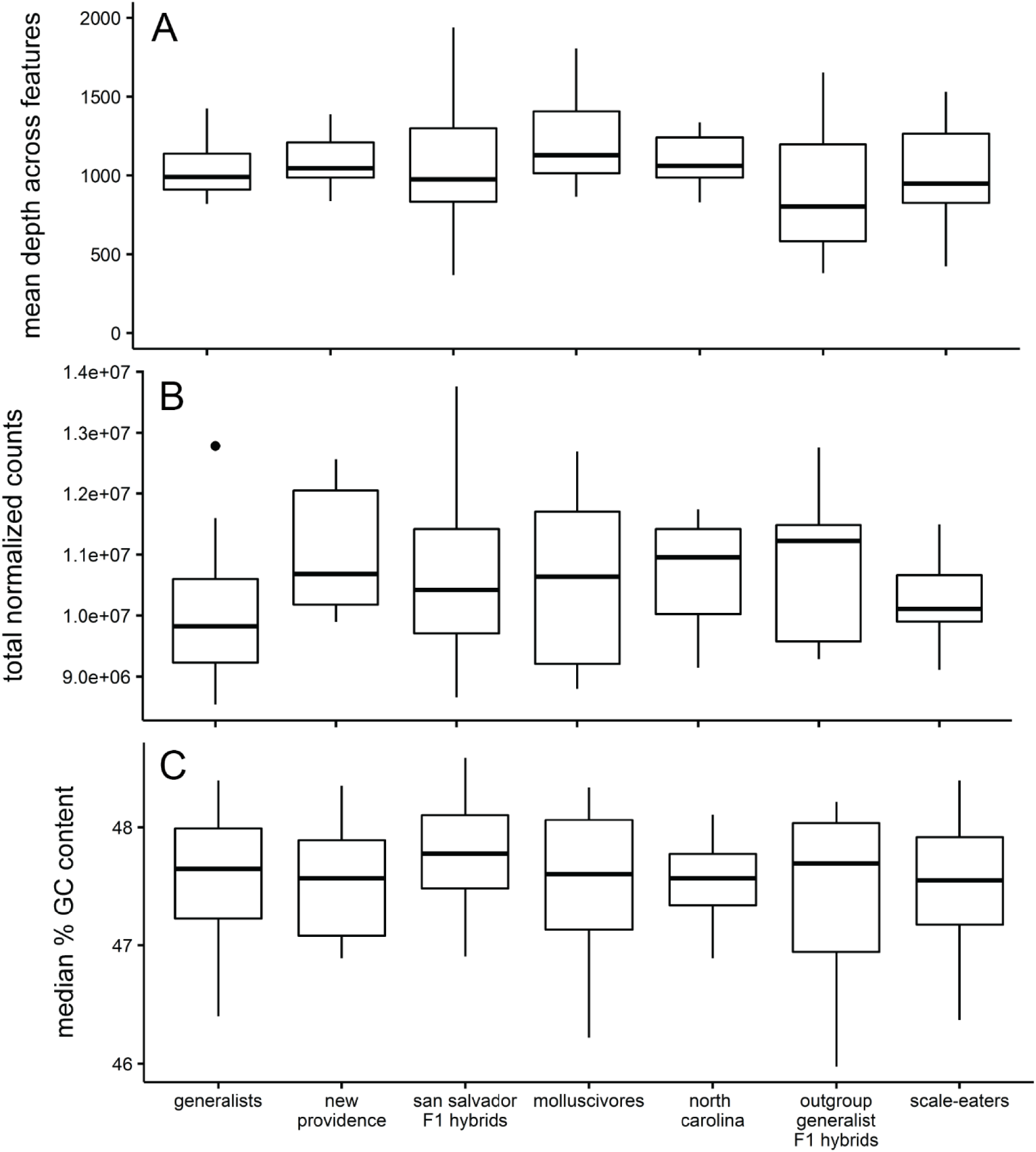
No significant difference among F1 purebred and F1 hybrid samples for A) mean read depth across annotated features (ANOVA; *P* = 0.32), B) total normalized read counts (ANOVA; *P* = 0.16), C) median percent GC content of reads (ANOVA; *P* = 0.32).

**Fig S8.**
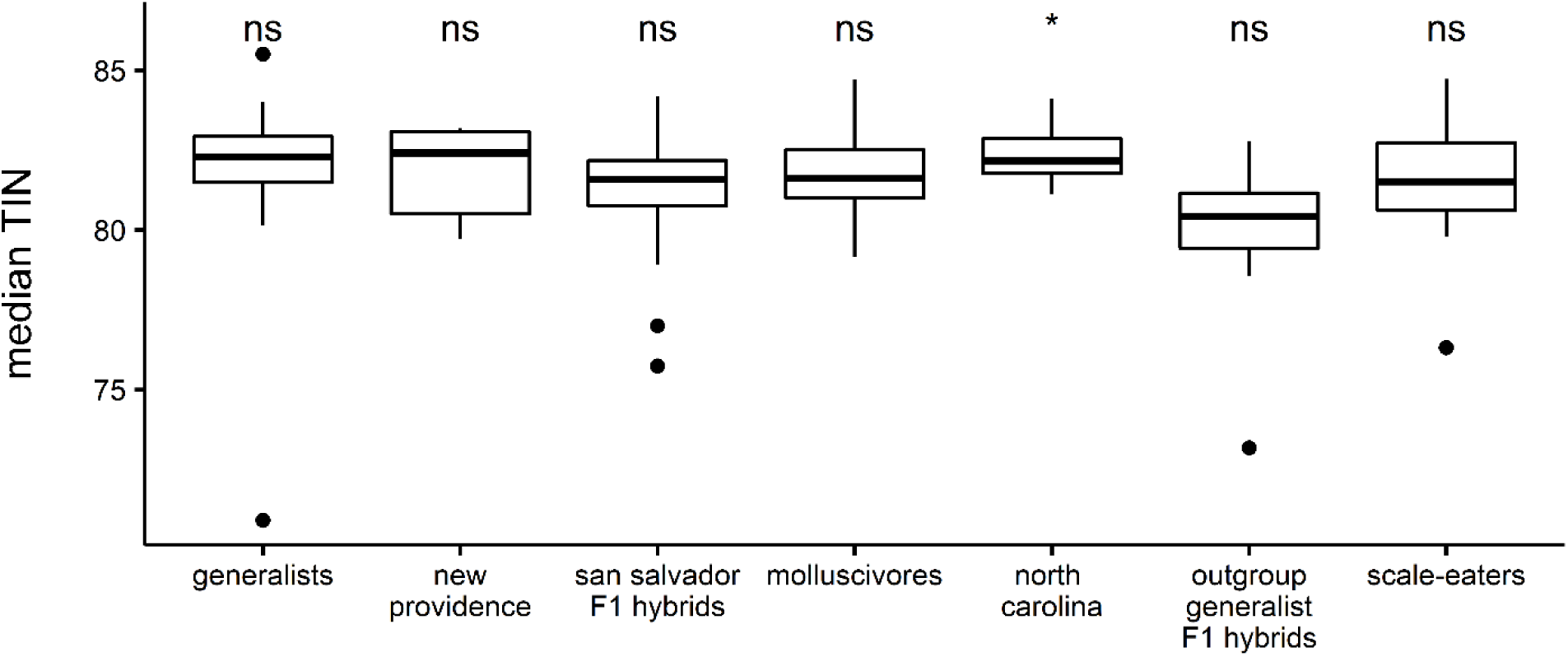
Median transcript integrity numbers for each species and generalist population. Tukey post-hoc test: *P* < 0.05 = *; *P* > 0.05 = ns.

**Fig S9.**
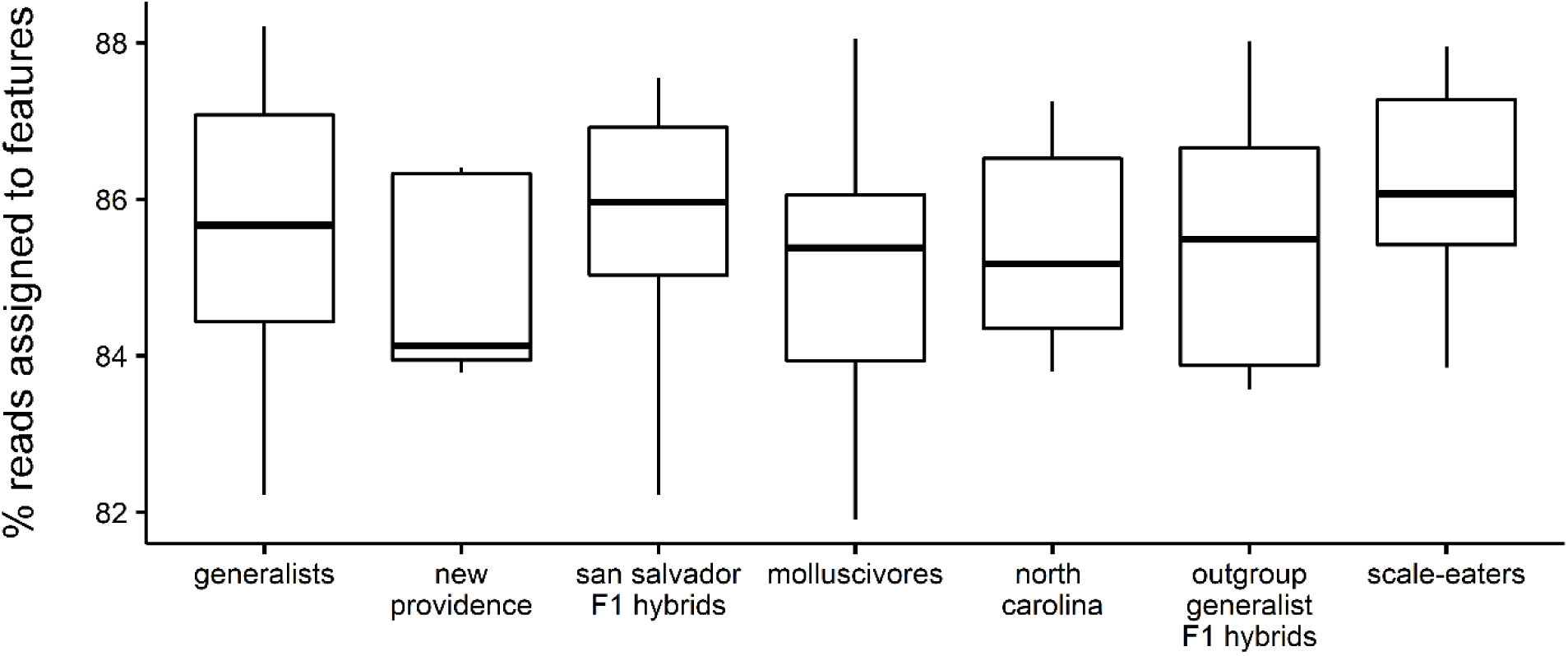
No significant difference in the percentage of reads mapping to annotated features of the *Cyprinodon* reference genome among F1 purebred and F1 hybrid samples (ANOVA; *P* = 0.17).

**Fig S10.**
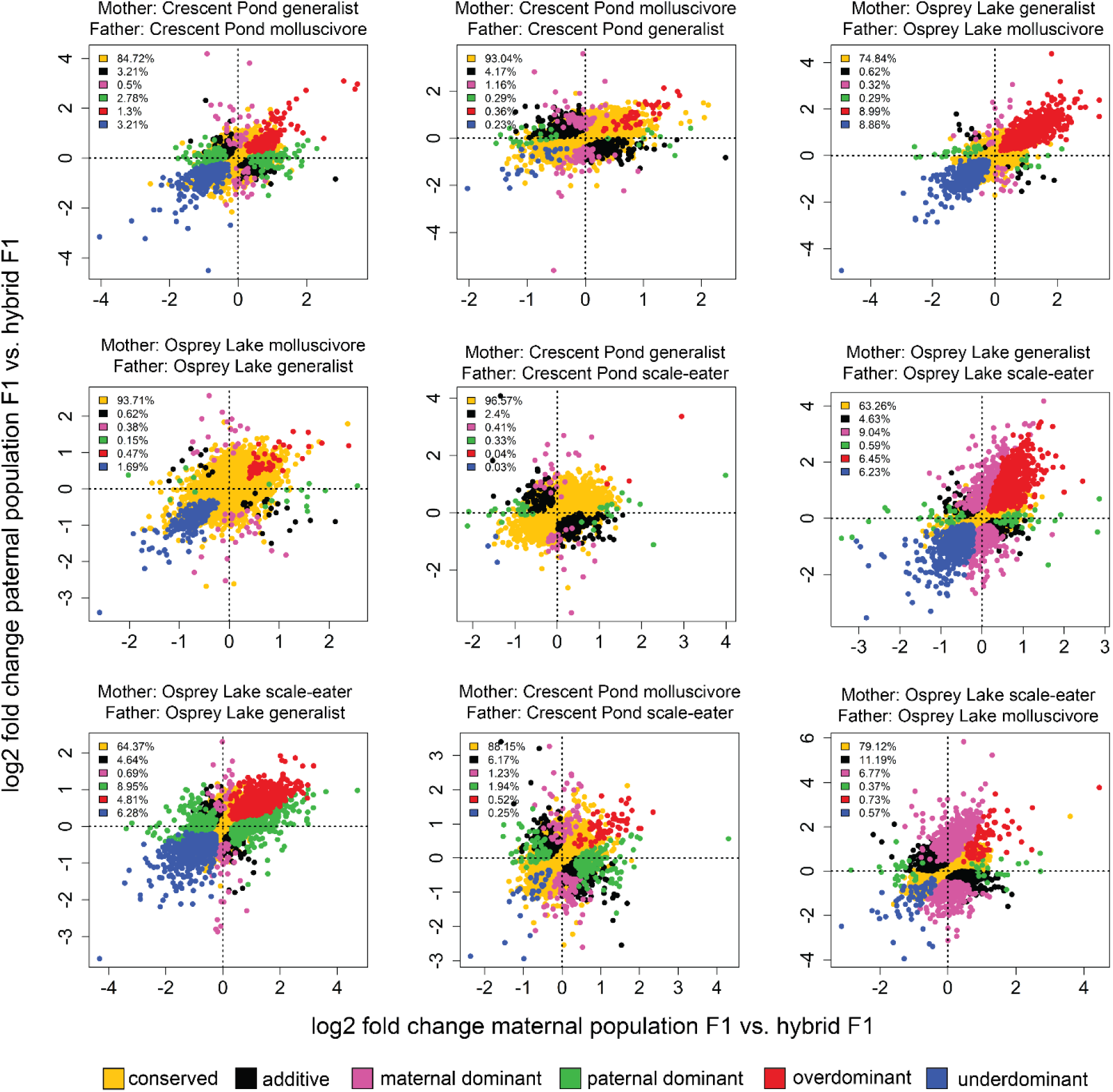
Gene expression inheritance for 2 dpf San Salvador hybrid crosses. Yellow = conserved (no difference in expression between groups or ambiguous expression patterns), black = additive (differential expression between purebred F1 and intermediate expression levels in hybrid F1), pink = maternal dominant (differential expression between purebred F1, differential expression between paternal population purebred F1 and F1 hybrids, no differential expression between maternal population purebred F1 and F1 hybrids), green = paternal dominant (differential expression between purebred F1, differential expression between maternal population purebred F1 and F1 hybrids, no differential expression between paternal population purebred F1 and F1 hybrids), red = overdominant (F1 hybrid gene expression significantly higher than parental population purebred F1), blue = underdominant (F1 hybrid gene expression significantly lower than parental population purebred F1).

**Fig S11.**
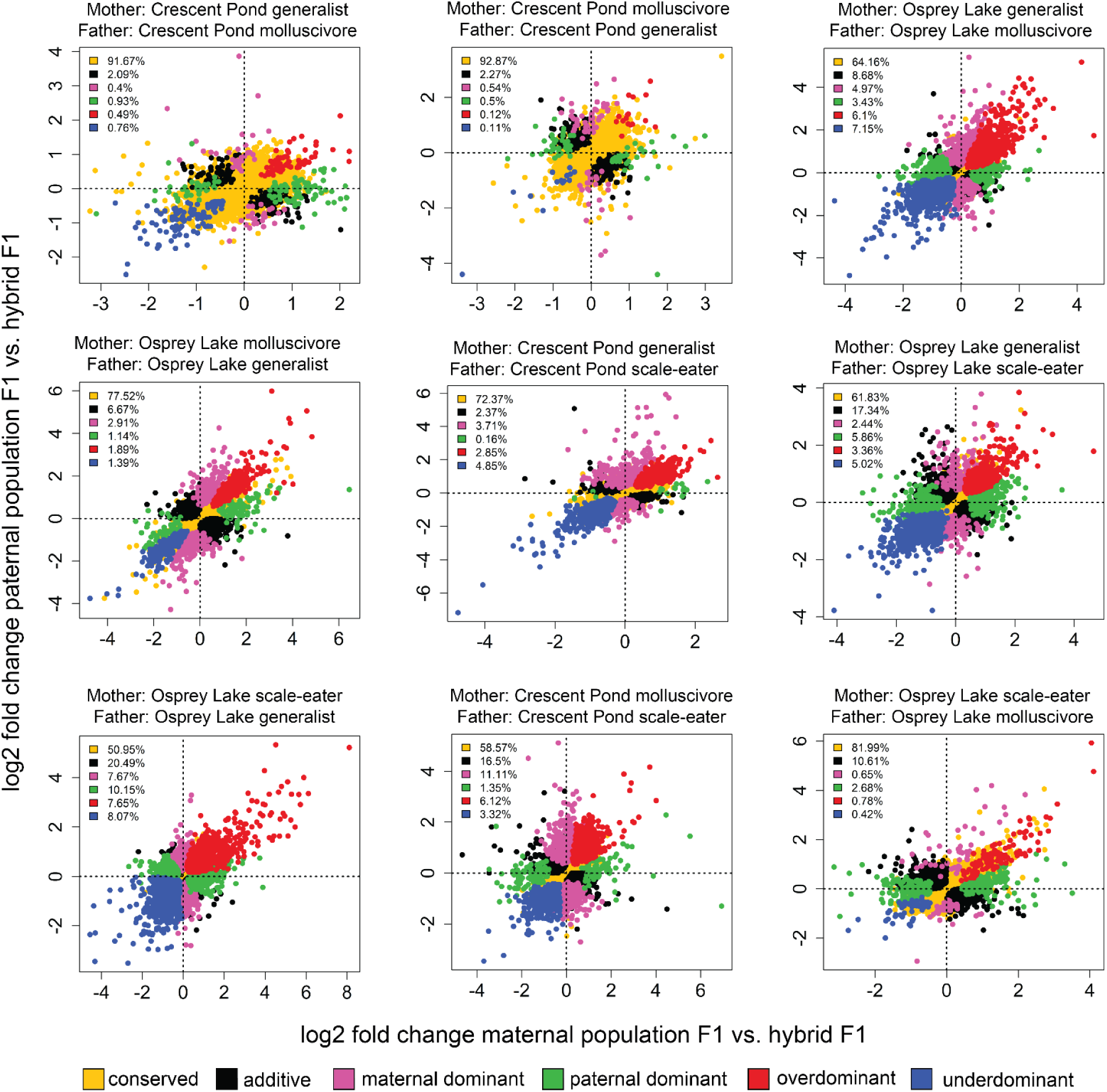
Gene expression inheritance for 8 dpf San Salvador hybrid crosses. Yellow = conserved (no difference in expression between groups or ambiguous expression patterns), black = additive (differential expression between purebred F1 and intermediate expression levels in hybrid F1), pink = maternal dominant (differential expression between purebred F1, differential expression between paternal population purebred F1 and F1 hybrids, no differential expression between maternal population purebred F1 and F1 hybrids), green = paternal dominant (differential expression between purebred F1, differential expression between maternal population purebred F1 and F1 hybrids, no differential expression between paternal population purebred F1 and F1 hybrids), red = overdominant (F1 hybrid gene expression significantly higher than parental population purebred F1), blue = underdominant (F1 hybrid gene expression significantly lower than parental population purebred F1).

**Fig S12.**
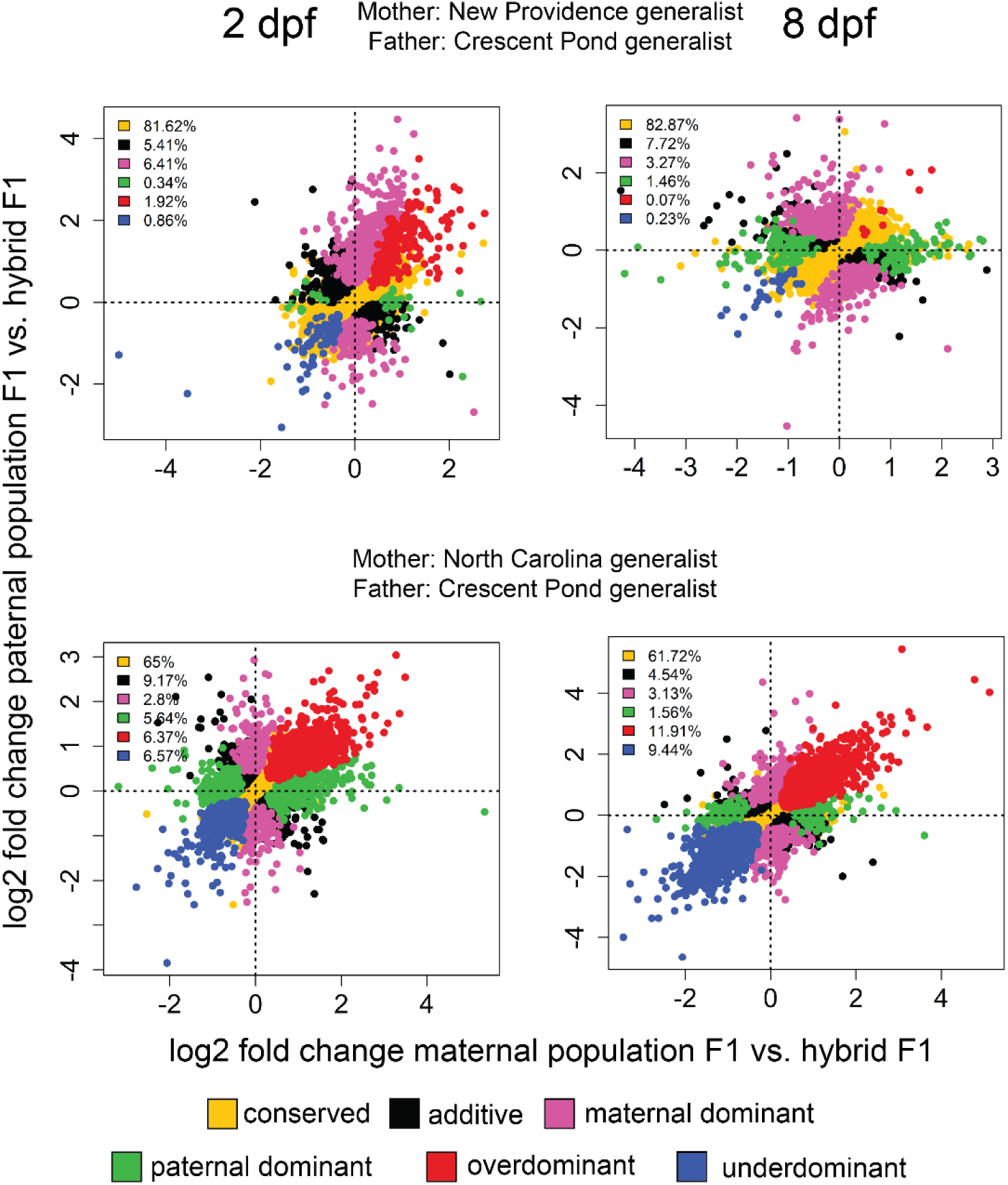
Gene expression inheritance for outgroup generalist population hybrid crosses. Yellow = conserved (no difference in expression between groups or ambiguous expression patterns), black = additive (differential expression between purebred F1 and intermediate expression levels in hybrid F1), pink = maternal dominant (differential expression between purebred F1, differential expression between paternal population purebred F1 and F1 hybrids, no differential expression between maternal population purebred F1 and F1 hybrids), green = paternal dominant (differential expression between purebred F1, differential expression between maternal population purebred F1 and F1 hybrids, no differential expression between paternal population purebred F1 and F1 hybrids), red = overdominant (F1 hybrid gene expression significantly higher than parental population purebred F1), blue = underdominant (F1 hybrid gene expression significantly lower than parental population purebred F1).

**Fig S13.**
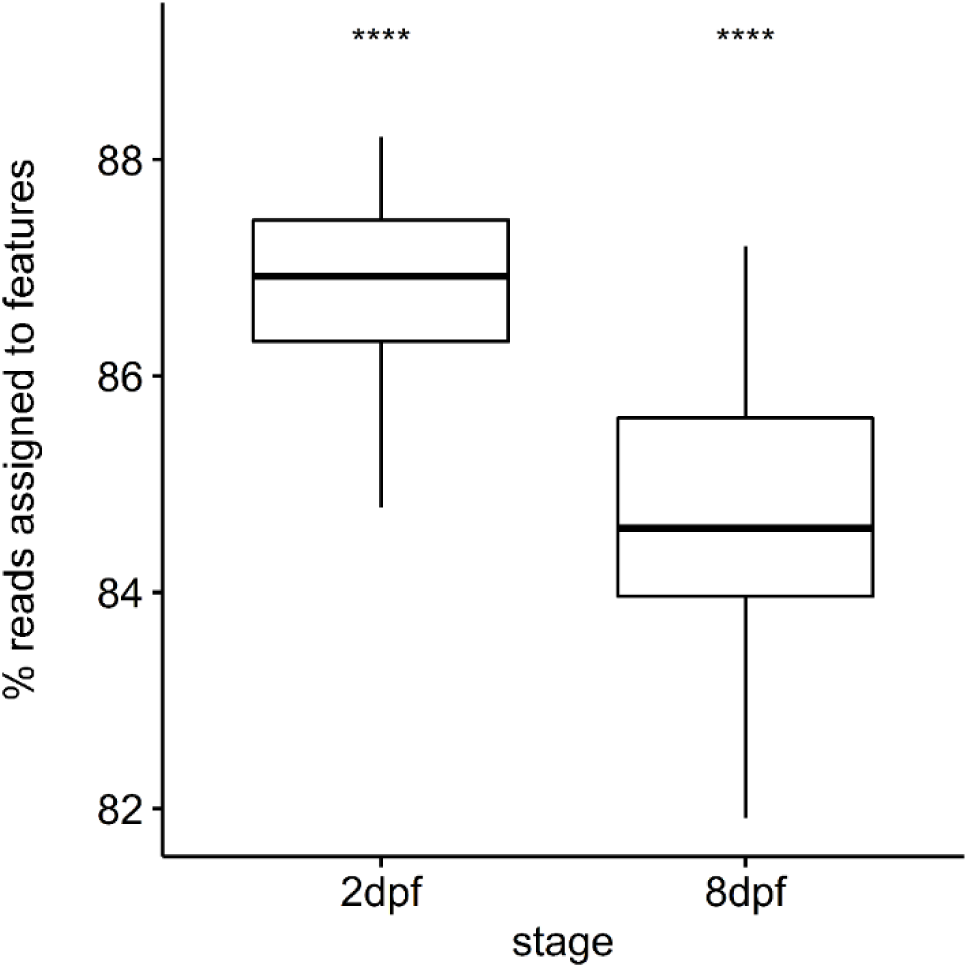
More reads assigned to features for 2 dpf samples than 8 dpf samples (Student’s *t*-test; P < 2.2 × 10^-16^).

**Fig S14.**
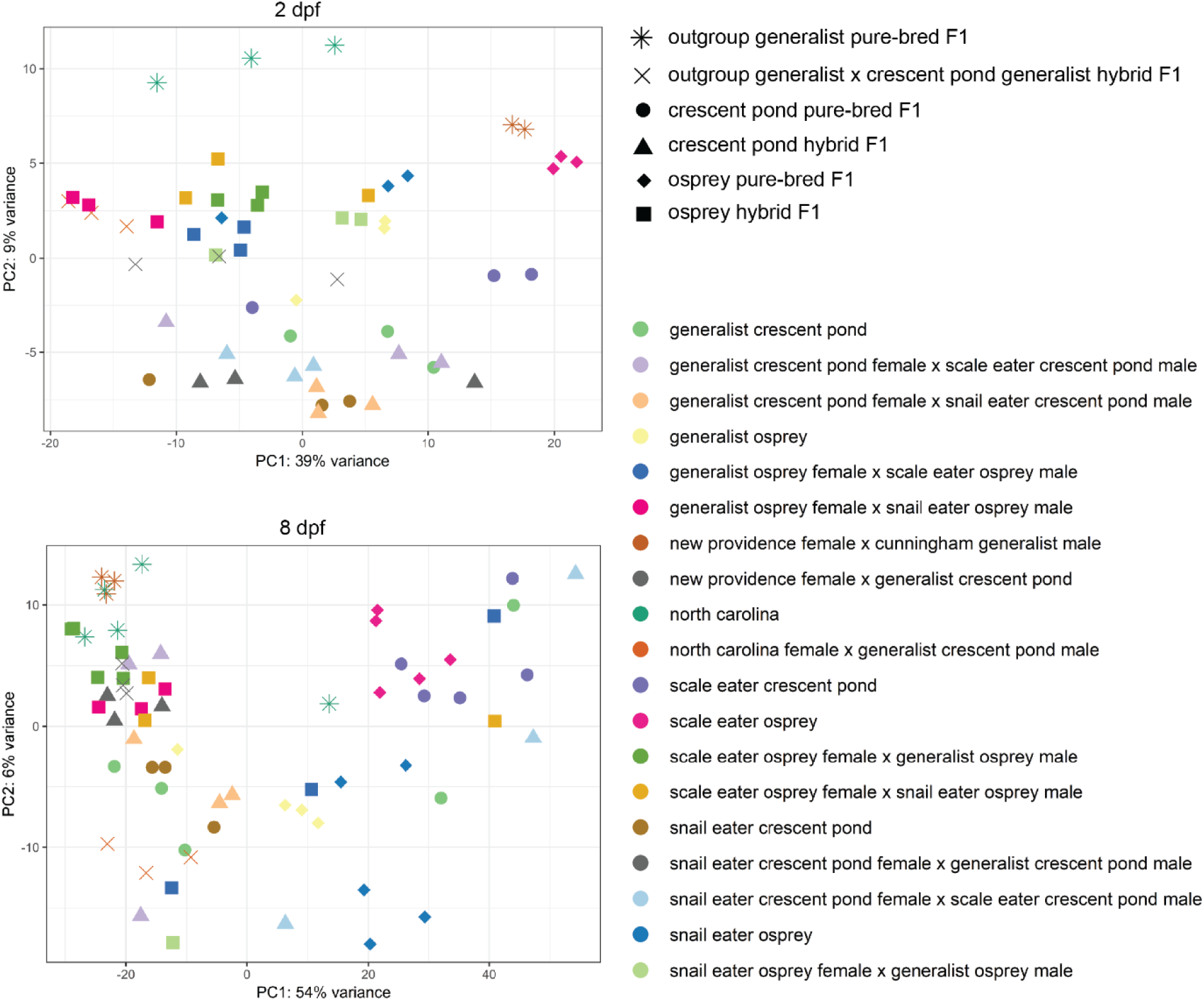
First two principal components explaining 48% (2 dpf) and 60% (8 dpf) of the variance across normalized read counts.

